# Hidden in the sand: phylogenomics unravel an unexpected evolutionary history on the desert-adapted vipers of the genus *Cerastes*

**DOI:** 10.1101/2023.10.03.560686

**Authors:** Gabriel Mochales-Riaño, Bernat Burriel-Carranza, Margarida Isabel Barros, Guillermo Velo-Antón, Adrián Talavera, Loukia Spilani, Héctor Tejero-Cicuéndez, Pierre-André Crochet, Alberto Piris, Luis García-Cardenete, Salem Busais, Johannes Els, Mohammed Shobrak, José Carlos Brito, Jiří Šmíd, Salvador Carranza, Fernando Martínez-Freiría

## Abstract

The desert vipers of the genus *Cerastes* are a small clade of medically important venomous snakes within the family Viperidae. According to published morphological and molecular studies, the group is comprised by four species: two morphologically similar and phylogenetically sister taxa, the African horned viper (*Cerastes cerastes*) and the Arabian horned viper (*Cerastes gasperettii*); a more distantly related species, the Saharan sand viper (*Cerastes vipera*), and the enigmatic Böhme’s sand viper (*Cerastes boehmei*), only known from a single specimen in captivity allegedly captured in Central Tunisia. In this study, we analyzed one mitochondrial marker (COI) as well as genome-wide data (ddRAD sequencing) from 28 and 41 samples, respectively, covering the entire distribution range of the genus to explore the population genomics, phylogenomic relationships and introgression patterns within the genus *Cerastes*. Additionally, and to provide insights into the mode of diversification of the group, we carried out niche overlap analyses considering climatic and habitat variables. Both nuclear phylogenomic reconstructions and population structure analyses have unveiled an unexpected evolutionary history for the genus *Cerastes*, which sharply contradicts the morphological similarities and previously published mitochondrial approaches. *Cerastes cerastes* and *C. vipera* are recovered as sister taxa whilst *C. gasperettii* is a sister taxon to the clade formed by these two species. We found a relatively high niche overlap (OI > 0.7) in both climatic and habitat variables between *C. cerastes* and *C. vipera*, contradicting a potential scenario of sympatric speciation. These results are in line with the introgression found between the northwestern African populations of *C. cerastes* and *C. vipera*. Finally, our genomic data confirms the existence of a lineage of *C. cerastes* in Arabia. All these results highlight the importance of genome-wide data over few genetic markers to study the evolutionary history of species.

## Introduction

Species-level phylogenies are crucial for understanding evolution and speciation (Barraclough & Nee, 2001). An accurate knowledge of phylogenetic relationships is critical to study multiple evolutionary processes such as interspecific diversification, trait evolution or adaptative responses of species to changing environments, among others (Guo et al., 2019; Schoch et al., 2009; Zhang et al., 2022). Traditionally, mitochondrial markers have been widely used in phylogenetic and/or phylogeographic approaches to infer the evolutionary history of species (Brown, 2002). However, mitonuclear discordances have been widely reported in many taxa (e.g., Dinis et al., 2019; Burriel-Carranza et al., 2023a; Hinojosa et al., 2019; Mochales-Riaño et al., 2023; Zaidi & Makova, 2019), mainly due to incomplete lineage sorting, sex-biased asymmetries, introgression, or different selection pressures on mitochondrial and nuclear genomes (Toews & Brelsford, 2012). Extensive discussion about this topic has led to the general agreement that the use of genomic data can efficiently resolve the evolutionary relationships of target species (e.g., Mao et al., 2019), as well as to infer other mechanisms, like introgression, which may influence the evolutionary trajectory of species (Cahill et al., 2015; Green et al., 2010). Remarkably, the advent of molecular genomics has allowed for the identification and characterization of introgression (i.e., the process by which genetic material from one species is incorporated into the genome of another species through interbreeding; Harrison & Larson, 2014) with increasing accuracy (Cahill et al., 2015; Leonard et al., 2013; Ottenburghs et al., 2017), confirming its significant impact on the evolution, adaptation and diversity of many species, including adaptive radiations (Seehausen, 2004). Adaptive introgression (the transfer of alleles from one species to another favored by natural selection) has even been proposed as another potential source of adaptation, aside from new mutations or standing variation (Hedrick, 2013), and has been reported to be involved in pesticide resistance, color polymorphism or adaptation to arid environments, among others (Anderson et al., 2009; Liu et al., 2021; Nanaei et al. 2023; Rocha et al. 2023; Song et al. 2011).

Arid areas encompass a large proportion of the Earth’s surface and are important for understanding global biodiversity patterns and especially patterns of reptile diversity (Brito et al., 2014; Roll et al., 2017; Tejero-Cicuéndez et al., 2022a). Arid areas are often inhabited by specialized arid-adapted species, showing high levels of inter- and intra-specific diversity and endemicity (e.g., Burriel-Carranza et al., 2023b; Garcia-Porta et al., 2017; Metallinou et al., 2012, 2015; Velo-Antón et al 2018). Much attention has been focused on arid areas from Australia and North America, while North Africa and Arabia have been traditionally neglected despite an increasing number of biodiversity studies carried out in the last decade (e.g., Brito et al., 2014; Carranza et al., 2018; Metallinou et al., 2012, 2015; Gonçalves et al., 2018a; Martínez-Freiría et al., 2017; Šmíd et al., 2021a; Tejero-Cicuéndez et al., 2022b; Burriel-Carranza et al., 2019, 2023a, 2023b). These studies have reported high levels of cryptic diversity in these relatively underexplored regions and have emphasized vicariant processes as the primary factor driving speciation/diversification. Additionally, they have provided valuable insights into how the onset and expansion of deserts, in conjunction with Plio-Pleistocene climatic cycles, have influenced the diversification processes of the region’s biota. However, there has been limited consideration of other evolutionary processes, such as ecological adaptation and lack of geographic isolation (as seen in sympatric speciation), which require further exploration in Palearctic deserts (Brito et al., 2014). In these arid regions most studies on reptiles have generally focused on lizards, while other groups, such as snakes, have been relatively neglected and under-explored (but see *Daboia* vipers, Martínez-Freiría et al., 2017; *Psammophis schokari*, Gonçalves et al., 2018a).

The desert vipers of the genus *Cerastes* (Family Viperidae, subfamily Viperinae) are a monophyletic group of medically important venomous snakes adapted to desert environments (Phelps, 2010; Alencar et al., 2016; Šmíd & Tolley, 2019). As other Palearctic vipers, they are highly dependent on climatic conditions and are susceptible to range shifts and demographic changes driven by climatic oscillations (Brito et al., 2011; Martínez-Freiría et al., 2015, 2017; Lucchini et al., 2023). The genus *Cerastes* is widely distributed across the arid regions of North Africa and the Middle East, particularly in the Sahara Desert and the Arabian Peninsula (Phelps, 2010; Sindaco et al., 2014). According to morphological (Wagner & Wilms, 2010; Werner et al., 1991), ecological (Brito et al., 2011) and phylogenetic studies, relying mostly on mtDNA data (Alencar et al 2016; Šmíd and Tolley, 2019), the group is comprised by four species: (A) two medium-sized sister species with generalist habitat requirements, the African horned viper (*C. cerastes* Linnaeus, 1758) and the Arabian horned viper (*C. gasperettii* Leviton & Anderson, 1967). Both species exhibit mostly allopatric distributions, with *C. cerastes* primarily inhabiting the Saharan desert and having a reported isolated population in southwestern Arabia, while *C. gasperettii* is exclusively found in arid regions within Arabia. Both species display individuals with and without horns. (B) A more distantly related species, the African sand viper *C. vipera* (Linnaeus, 1758), specialized in the sandy environments of the Sahara Desert. This species has dorsally positioned eyes and lacks horns entirely. A fourth species, Böhme’s sand viper (*Cerastes boehmei* Wagner & Wilms, 2010) is sometimes recognized (e.g., Uetz et al., 2023) but also disputed (e.g., Trape et al., 2023). It is based on a single specimen with aberrant horns, allegedly captured in Central Tunisia in 1991 and for which no genetic data was available so far. At the intraspecific level, molecular inferences have only been carried out in *C. gasperettii* (Carné et al., 2020), showing a low genetic variation. Morphological data has led to the description of several subspecies within the two horned vipers (i.e., *C. cerastes* and *C. gasperettii*): *C. c. cerastes* (widely distributed from Western Sahara to Egypt) and *C. c. hoofieni* (in southwestern Arabia based only on morphological data; Werner et al., 1991) within *C. cerastes*; and *C. g. gasperettii* (widely distributed in Arabia) and *C. g. mendelssohni* (Arava Valley; Werner et al., 1999) within *C. gasperettii*. A genomic study at the genus level would allow addressing the intraspecific variability and systematic of this group across North African and Arabia. Moreover, the combination of genomic and ecological data would help interpreting the evolutionary history of the group and enhance our understanding of its evolution and speciation history.

In this study, we used mitochondrial and double digest restriction-site associated DNA (ddRAD) sequencing data to infer the evolutionary history within the genus *Cerastes* and gain new insights into the intraspecific variability of *C. cerastes*. Given the remarkable patterns of morphological differentiation and habitat specialization in *Cerastes* species, we also implemented niche overlap analyses to explore the potential role of ecological adaptation in a putative process of sympatric speciation (e.g., Moutinho et al., 2020). Finally, following the niche overlap and FineRADstructure results, we tested for ancestral introgression between the three species. Our main objectives were: (1) to reassess the phylogenetic relationships within the genus *Cerastes* using novel genome-wide data, (2) to discuss possible drivers of speciation within the genus *Cerastes* across North Africa and Arabia and (3) to clarify the taxonomic status of *C. cerastes*.

## Materials and Methods

### Sample collection

A total of 36 samples representing the distribution range of *C. cerastes* (including *C. c. cerastes* and *C. c. hoofieni*)*, C. gasperettii* (*C. g. gasperettii*) and *C. vipera* were included in this study (Fig. 1, Fig. S1 and Table S1). No samples were available for *C. boehmei* and *C. g. mendelssohni*. Moreover, we included five individuals of *Echis omanensis* as outgroup (Šmíd & Tolley, 2019). Samples consisted of tail tips or scale clips in live specimens (n = 37), small fragments of body skin and muscle in roadkill specimens (n = 1) or stored in museum collections (n = 3) and were kept in 99% ethanol and stored at -20 °C upon arrival. For DNA extraction, samples were extracted using the MagAttract HMW Kit (Qiagen) following the manufacturer’s protocols.

**Fig. 1:**
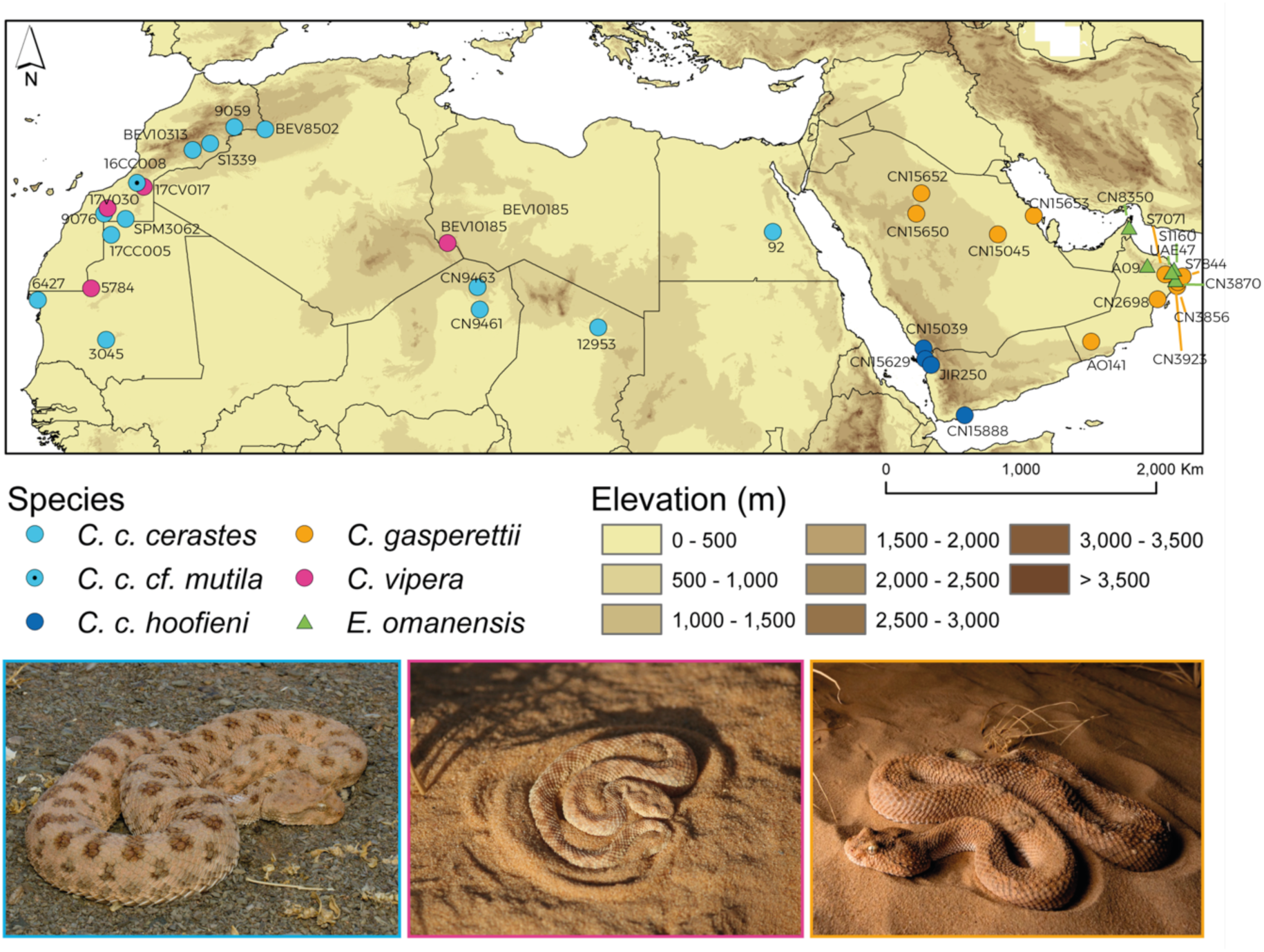
On the top, distribution of the individuals sampled in the study, colored per species or subspecies. On the bottom, pictures of the three species considered in this study, from left to right: *C. cerastes* (Morocco), *C. vipera* (Mauritania) and *C. gasperettii* (United Arab Emirates). Authors: F. Martínez-Freiría and A. Talavera.

### ddRADseq library preparation

We followed Peterson et al., (2012) protocol for double digest restriction site-associated DNA (ddRADseq) on a total of 41 samples. We double digested 500 ng of genomic DNA using a combination of a rare and a common cutting restriction enzymes (Sbf1 and Msp1, respectively). Fragments were purified with Agencourt AMPure beads before ligation with eight different barcoded Illumina adapters. Samples with unique adapters were then pooled, and each pool of eight samples was size-selected for fragments sized between 415 and 515 base pairs (bp). Illumina multiplexing indices were ligated to individual samples using a Phusion polymerase kit (high-fidelity Taq polymerase, New England Biolabs). Final pools were single-end sequenced on four lanes to a length of 75 base pairs (bp) Illumina NextSeq 500, at the UPF Genomics Core Facility (Barcelona, Spain).

### ddRADseq data processing

Raw reads were visually explored with FastQC (Andrews, 2010) to ensure data quality and absence of adapters. Then, reads for each individual were demultiplexed and processed with ipyrad v.0.9.84 (Eaton & Overcast, 2020), discarding sites with Phred score < 33, and reads with more than three missing sites. Consensus sequences with less than six reads, excessive heterozygous sites (more than three), or more than two alleles were also discarded. We created three different datasets: one to explore population structure analyses and gene flow containing all individuals of *Cerastes* together with *E. omanensis* (as outgroup), a second to explore more in-depth population structure analyses at the genus level containing only the genus *Cerastes* samples and a third dataset to explore intraspecific relationships with only *C. cerastes* samples. Then, for each dataset three independent ipyrad runs were performed using different clustering parameters for both filtered and consensus sequences (step 3 and 6 in ipyrad, respectively): 0.85, 0.89 and 0.92, selecting the clustering that maximized the number of samples without significantly reducing the number of loci retained (0.85 with the outgroup and 0.89 for both datasets without the outgroup). Then, following recommendations from O’Leary et al., (2018), we used Radiator (Gosselin et al., 2017), Plink2 (Chang et al., 2015) and VCFr (Knaus & Grünwald, 2017) implemented in a custom script (https://github.com/BernatBurriel/Post_processing_filtering) from Burriel-Carranza et al., (2023a) to filter iteratively and alternatively low-quality samples and loci. For all datasets, values of missing data allowance ranged from 98% to 60% of missing genotype call rate and missing data per individual, decreasing 2% between iterations. Subsequently, we applied a hard threshold of missing genotype call rate of 60%. We removed non-biallelic SNPs, applied a minor allele frequency filter (maf < 0.05) and removed monomorphic sites. For the different analyses, we retrieved *loci*, *haplotype* and *SNP* data (with or without an outgroup, depending on the analyses’ requirements): *loci* and *haplotype* datasets were generated with a second round of ipyrad after removing all individuals that did not pass the previously explained iterative filters and retaining only loci that were at least in 60% of all individuals. The *SNP* dataset (with the outgroup) was used for the introgression analysis, where all previously filtered SNPs were kept. For population structure analyses, we filtered the *SNP* dataset to only keep unlinked SNPs, selecting the SNP with the highest depth for each locus (from now on *uSNP* dataset). Finally, the *SNP* dataset only containing *C. cerastes* individuals was used for a Mantel test.

### Population structure analyses

We performed a Principal Component Analysis (PCA) with the unlinked *SNP* dataset (a total of 2,856 *uSNPS*) containing only *Cerastes* samples using Plink v1.9 (Chang et al., 2015). Then, we used ADMIXTURE (Alexander et al., 2009) to analyze the same dataset to detect population ancestry from K=2 to K=5. A total of 20 replicates for each K were calculated, selecting the best K value after 20 cross validations. We also used fineRADstructure (Malinsky et al., 2018) on the *haplotype* dataset to explore population structure within the three species, plus the outgroup, setting the minimum number of samples per loci to four. This analysis is optimal to detect different levels of structure within and between populations and to explore co-ancestry patterns for non-model organisms (Malinsky et al., 2018). We also performed a Mantel test with the *Cerastes cerastes* samples (*SNP* dataset containing only *C. cerastes* samples). Lastly, we calculated genetic diversity and pairwise Fst values with GenoDive v.3.0 (Meirmans, 2020) with 10,000 permutations at the species and subspecies level, using the *SNP* dataset. Visualization of results from these analyses was performed with R v.3.6.3 (R Core Team, 2021) and the R package ggplot2 (Wickham, 2016).

### COI sequencing and phylogenetic inference

We sequenced a COI fragment for a total of 24 individuals for the three species within the *Cerastes* genus (information regarding the specimens used can be found in Table S1). Primers and PCR amplification conditions were implemented according to Nagy et al., (2012). PCR products were purified and Sanger sequenced by Genewiz (UK). Then, we performed a multiple sequence alignment with MAFFT (Katoh & Standley, 2013) for the 24 samples together with two previously sequenced samples (13693 and 12953, accession number ON943580 and ON943571, respectively, from Velo- Antón et al., 2022) as well as two outgroups (two individuals of *Echis carinatus*, accession numbers MG699965 and MG699966, respectively). We used BEAST2 v.2.6.4 (Bouckaert et al., 2019) to reconstruct a Bayesian Inference (BI) time-calibrated tree. We calibrated the phylogeny dating the deepest node in the phylogeny (between *Echis* and *Cerastes*), as suggested by Stange et al., (2018) and also implemented in previous studies focused on reptiles (Burriel-Carranza et al., 2023a, 2023b; Thanou et al. 2023), with a normal distribution from a mean age of 28 million years ago (Mya) (Šmíd & Tolley, 2019) and encompassing a 95% HPD interval (25.65-31.55 Mya). We selected a GTR model with four gamma categories (base frequencies and proportion of invariant sites were estimated), and a relaxed clock LogNormal was used with a coalescent constant population tree prior. We conducted four independent runs of 10^9^ generations sampling every 10^4^ generations. Convergence was checked with Tracer v.1.7.2 (Rambaut et al., 2018). Posterior distributions were combined with LogCombiner v.2.6.3, discarding 10% of the posterior trees as burn-in and a maximum clade credibility tree was obtained calculating median heights in TreeAnnotator v.2.6.3 (BEAST2 v.2.6.4; Bouckaert et al., 2019).

### Phylogenomic reconstructions

We reconstructed the phylogenomic history of the three species of *Cerastes* using three different phylogenetic methods. First, with the concatenated *loci* dataset (531,260 bp, including the outgroup) we reconstructed a Maximum Likelihood (ML) tree. For that, we used RaxML-NG (Kozlov et al., 2019), with the GTR+I+G model with a total of 100 starting trees (50 random and 50 parsimony) and performing 1,000 bootstraps for branch support. Later, we filtered the *loci* dataset to maintain only loci without missing data, obtaining a total of 627 loci and 40,831 bp (including the outgroup). With those, we used BEAST2 v.2.6.4 (Bouckaert et al., 2019) to obtain a time-calibrated tree. We used the same calibration point as described for the mitochondrial phylogeny. We selected a GTR model with four gamma categories (base frequencies and proportion of invariant sites were estimated), and a relaxed clock LogNormal was used with a coalescent exponential population tree prior. We conducted four independent runs of 10^8^ generations sampling every 10^4^ generations. Posterior distributions were combined with LogCombiner v.2.6.3. Convergence was assessed with Tracer v.1.7.2 (Rambaut et al., 2018), applying a 10% burn-in. Finally, and to confirm the unexpected phylogenomic relationships (see results), we calculated a species tree using SNAPP (Bryant et al., 2012). This method allows each SNP to have its own history under the multispecies coalescence model, while bypassing the need to sample individual gene trees (Bryant et al., 2012). Due to computational constraints, we ran an extra ipyrad selecting only two individuals per species (Table S1) and obtaining a total of 6,668 SNPs (including the outgroup), allowing a total of 20% of missing data with each SNP present in at least one individual per species. We used the same calibration point as described for the mitochondrial phylogeny. We ran four different SNAPP analyses for 1,000,000 Markov chain Monte Carlo (MCMC) generations, sampling every 1,000 steps, setting mutation rates (*u* and *v*) to 1.0 and using different gamma prior distributions for alpha and beta (2, 200; 2, 2,000; 2, 20,000). Posterior distributions were combined with LogCombiner v.2.6.3. Convergence to stationary distributions was observed after 10% generations, and hence 10% of posterior trees were removed as burn-in with Tracer v.1.7.2 (Rambaut et al., 2018) and the posterior distribution was considered adequately sampled with an effective sample size higher than 200 for all parameters. Trees were combined with LogCombiner v.2.6.3 and visualized in DensiTree v2.2.6 (Bouckaert, 2010). Alternative prior combinations produced highly concordant results. We produced a second species tree under the same conditions but including two specimens of *C. c. hoofieni* to the same dataset (obtaining 6,379 SNPs) to study possible different evolutionary histories of the two *C. cerastes* subspecies.

### Niche overlap analyses

In order to investigate the potential role of environmental adaptation in the diversification of the genus, we performed pairwise niche overlap tests between the major clades within the genus, resulting from our ddRADseq inferences, considering two geographic scales and two types of variables. The global scale analyses included a total of 990 records at a spatial resolution of 5 arc minutes (∼10x10km; 561 *C. cerastes*, 213 *C. gasperettii* and 216 *C. vipera*) spanning across the whole distribution of the three species (Fig. S2), and two distinct sets of low correlated variables (R < 0.72; Table S2): (1) a climatic dataset, which included seven precipitation- and temperature-related variables downloaded from WorldClim v.2.1 (www.worldclim.org), and (2) a landcover dataset, which comprised three percentages of landcover variables, which resulted from calculating the number of 10 arc seconds resolution pixels of three landcover categories representative of the Saharan and Arabian deserts (from the ESA GlobCover ver 2.2, Bicheron et al., 2008) in 5 arc minutes resolution pixels, using the function aggregate from ArcGIS v.10.5 (ESRI, 2016). Occurrence data for the global scale analysis were gathered from distinct sources, including fieldwork conducted by the authors for the period 2001-2021 (n = 241), bibliographic references (n = 180; e.g., Werner et al., 1999; Brito et al., 2011; García- Cardenete et al., 2017), museum collections (n = 106; e.g., NHM-London, MNHN-Paris) and GBIF (n = 463; www.gbif.org, 2021). Due to the underrepresentation of the landcover heterogeneity for the Saharan and Arabian deserts as available in the ESA GlobCover dataset, we also approached niche overlap for the Saharan species at a regional scale of the Atlantic Sahara (i.e., Mauritania and Western Sahara), where a detailed landcover map was recently derived (Campos et al., 2018). Consequently, for this regional scale analysis, we considered a subset of 170 records (118 for *C. cerastes* and 52 for *C. vipera*), available at a higher accuracy (i.e., 30 arc seconds, ∼1x1km), since 69% were gathered during fieldwork campaigns (the remaining from bibliography, 19% and GBIF, 12%). In addition, we used seven percentages of landcover variables (see Table S2), which resulted from using the function aggregate from ArcGIS over seven landcover categories or groups of categories with 1 arc second resolution pixel (see Table S2) to new rasters with 30 arc seconds resolution pixel.

For the three approaches (global-climate, global-landcover and regional-landcover), occurrence data were used to extract the corresponding variable values and then run three distinct PCA. Following a three-dimensional hypervolume approach, the first three components of each PCA were used to describe the environmental niche of each major clade (i.e., total volume and unique part, intersection and union with the other species’ volume) as obtained from ddRAD data analyses and perform pairwise niche-overlap tests between them (see Martínez-Freiría et al., 2020 for a similar procedure). Analyses were performed in R with the ‘hypervolume’ package (Blonder et al., 2014), using a Silverman bandwidth estimator and a set of 999 random points to sample the kernel density to build hypervolumes. To quantify environmental niche overlap, the Sørenson index (K) and the Overlap Index (OI) were used (e.g., Martínez-Freiría et al., 2020). These statistics quantify predicted niche similarity, and they both range from 0 (no overlap) to 1 (identical niche). The Overlap Index (OI) relates the observed and maximum values of K, and is preferred when niches present different sizes (Simó-Riudalbas et al., 2018). In order to test the significance of the overlap, a randomization procedure similar to that applied by Warren et al., (2010) was used. By comparing the produced Ks and OIs, we tested whether the observed niche overlap was more different than the overlap that would be obtained by merging occurrences randomly. This procedure was performed 999 times to generate the null distribution, and to evaluate the significance of each overlap.

### Introgression analyses

To study ancestral introgression, we ran the ABBA-BABA test implemented in Dsuite (Malinsky et al., 2021) with the *SNP* dataset (32,254 SNPs), including the outgroup. To do so, we split our samples per species (and subspecies when possible) and calculated the D-statistics using the new phylogenetic topology. Patterson’s D-statistic uses asymmetry in gene tree topologies to quantify introgression between either of two lineages which share a common ancestor (P1 and P2) and a third lineage (P3) that diverged from the common ancestor of P1 and P2. A fourth individual (P4) is needed to inform the ancestral state for each comparison (Malinsky et al., 2021). In here, we set *E. omanensis* as P4 to set the ancestral state. *C. gasperetti* or *C. vipera* were used as P3 while *C. c. cerastes*, *C. c. hoofieni* or *C. vipera* were set as P1 and P2 terminals, depending on the comparison. Finally, we calculated the F- branch statistics to infer the excess of shared alleles between the groups, using *E. omanensis* as outgroup.

## Results

### ddRADseq data processing

Genome-wide data was sequenced for a total of 41 samples, including five individuals of *E. omanensis* from regions spanning Western Sahara to Oman (Fig. 1, Fig. S1 and Table S1). However, after the iterative filtering and due to the low DNA quality of some samples, we kept a total of 28 high-quality samples: four *C. vipera*, seven *C. gasperettii*, thirteen *C. cerastes* (seven *C. c. cerastes* and six *C. c. hoofieni*) and four *Echis omanensis* (Fig. S1 and Table S1). Total reads obtained from sequencing added up to 7.7 x 10^7^. After applying quality filtering, more than 99% of the raw reads remained, with an average of 2.7 x 10^6^ reads per individual (Table S1).

### Population structure analyses

Population genomic results of the unlinked *SNP* dataset (2,856 *uSNPs*) showed an unexpected result (Fig. 2). PC1 split *C. gasperettii* from both *C. cerastes* and *C. vipera* individuals whilst PC2 separated *C. cerastes* from *C. vipera* (Fig. 2A). The *C. c. hoofieni* individuals clustered together with the rest of African *C. cerastes* (Fig. 2A). PC3 sorted *C. cerastes* following their geographical locations (northwestern Africa, northeastern Africa and Arabia). The best supported scenario in Admixture recovered the three different species for K=3 (Fig. 2B. For K=4, the samples of *C. c. hoofieni* formed a unique cluster, with the two geographically closer African individuals (92 and 12953; Table S1) sharing a limited amount of coancestry (Fig. 2B). FineRADstructure analyses showed five different clusters (including the outgroup) representing the different species and with *C. c. hoofieni* having a different cluster (Fig. 2C). Again, *C. gasperettii* was the most differentiated within the *Cerastes* genus and showed high similarity among samples. *Cerastes vipera* also displayed a high similarity among samples, with only the individual from central North Africa being less similar than the rest. *Cerastes cerastes* individuals showed three main clusters corresponding with their geographical locations (northwestern Africa, northeastern Africa and Arabia). *Cerastes vipera* samples reported higher shared co-ancestry with the African samples of *C. c. cerastes* than with the Arabian individuals of *C. c. hoofieni* (Fig. 2C). Mantel test analysis showed strong isolation by distance among *C. cerastes* individuals (r = 0.69, *p*-value < 0.001) (Fig. S3). The Fst comparisons revealed consistently high values in all pairwise comparisons, indicating a strong isolation among species (Table S3), being similar between *C. c. cerastes* and both *C. c. hoofieni* and *C. vipera*. Finally, genetic diversity measures consistently indicated lower observed diversity compared to the expected values (Table S4), with *C. c. hoofieni* displaying the lowest levels of heterozygosity.

**Fig. 2:**
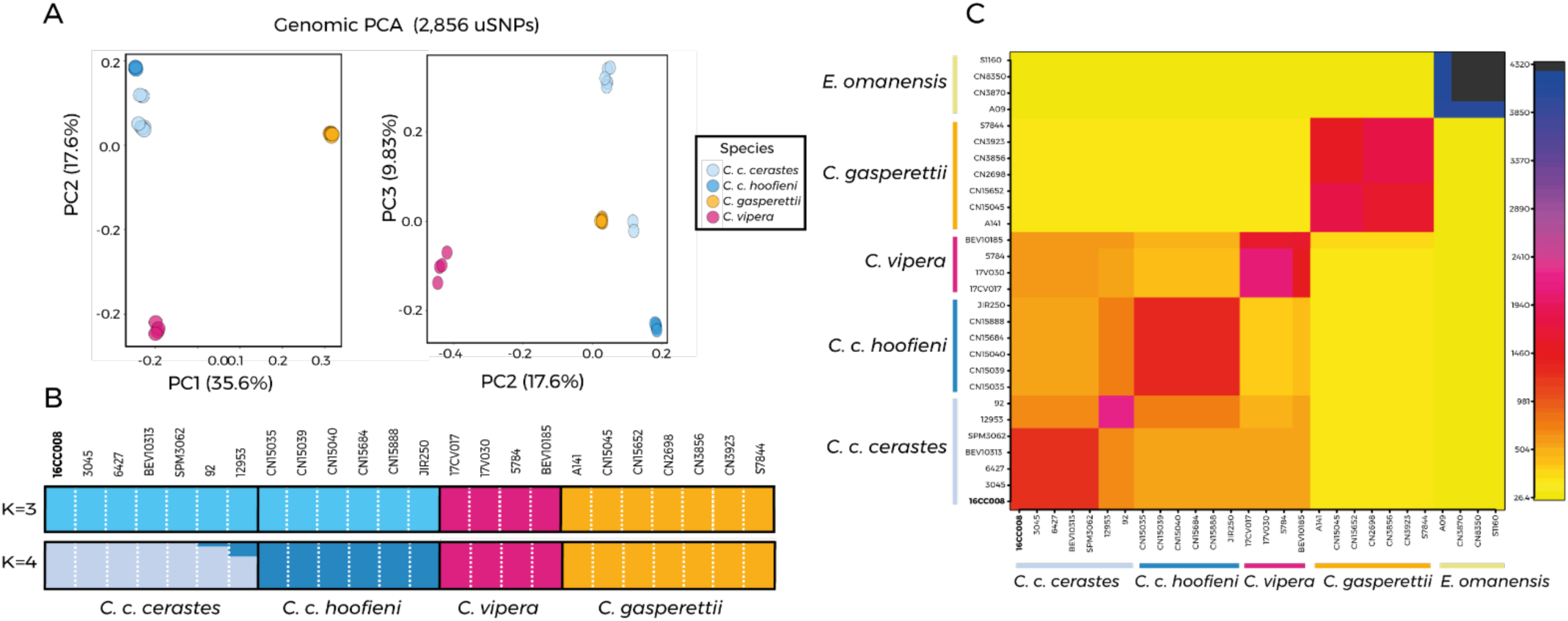
Population structure analyses for the three species of *Cerastes*, with sample colors per species or subspecies. A) Principal Component Analysis (PCA) with a total of 2,856 *uSNPs*. B) Admixture analyses at the individual level for clusters with K=3 and K=4. C) Co-ancestry matrix from fineRADstructure, showing pairwise similarity. Darker colors show higher shared co-ancestry. In bold the putative *C. c. mutila* individual (16CC008).

6.48

### Mitochondrial phylogenetics

The phylogenetic reconstruction of the mitochondrial COI gene fragment split the three different *Cerastes* species (Fig. 3A). *Cerastes vipera* represented the deepest split within the genus, dating 12.78 ± 5.62 Mya, while *C. cerastes* and *C. gasperettii* were recovered as sister taxa, splitting 9.19 ± 4.25 Mya. Within *C. cerastes*, there were two lineage splitting 3.76 ± 2.6 Mya: a first one containing individuals from Morocco, Western Sahara, Mauritania and Algeria, and a second lineage with the individuals from Chad and Egypt together with the Arabian *C. c. hoofieni*. All individuals within *C. gasperettii* clustered together with an individual from southern Oman showing a slight separation. Individuals of *C. vipera* formed a separated lineage.

**Fig. 3:**
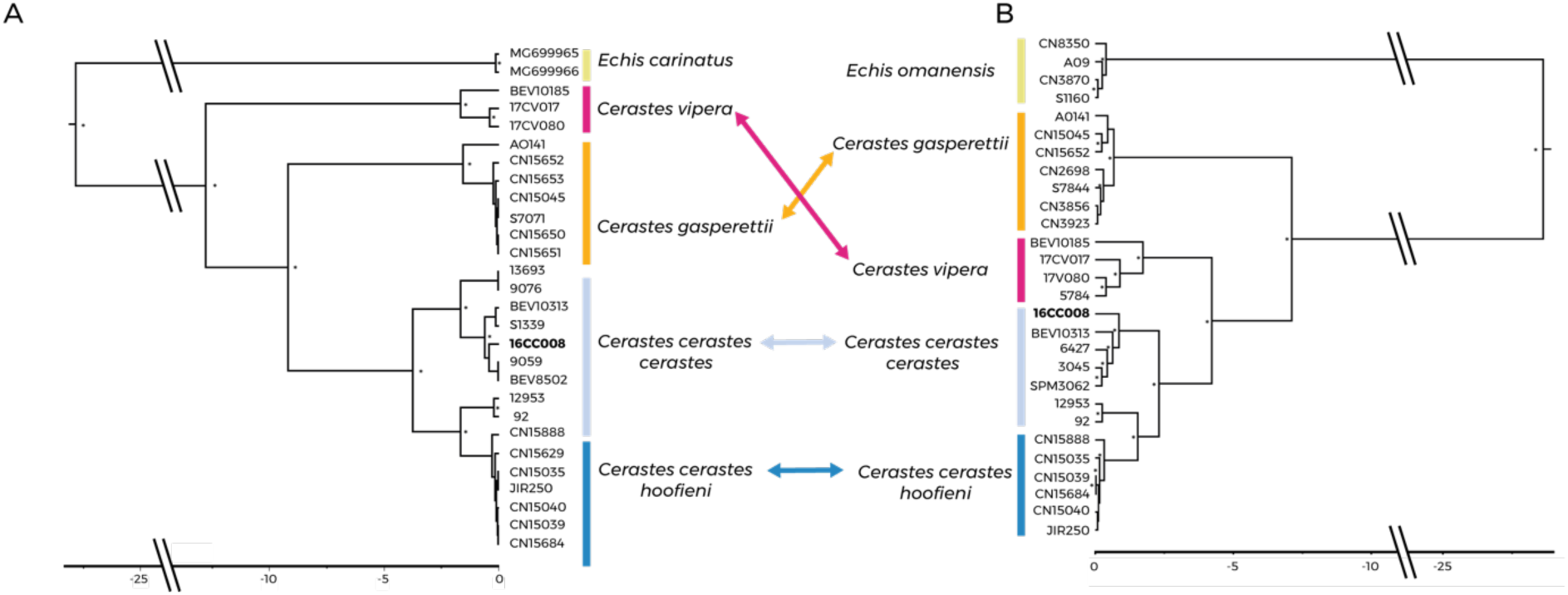
Mitonuclear discordance between mitochondrial and nuclear phylogenies. A) Mitochondrial Bayesian phylogeny for the COI gene for the species and subspecies included in the study. B) Bayesian time-calibrated tree produced with BEAST2 v.2.6.4 based on 627 concatenated loci and covering a total of 40,831 bp. Divergence time between *C. gasperettii* and the ancestor of both *C. cerastes* and *C. vipera* occurred around 7.12 ± 3.83 Mya and between *C. cerastes* and *C. vipera* around 4.22 ± 2.23 Mya. Posterior probabilities above 0.95 are indicated with an asterisk. In bold is shown the putative *C. c. mutila* sample.

### Phylogenomic reconstruction

The ML phylogenomic tree, with 531,260 bp, showed *C. vipera* and *C. cerastes* as sister species, whilst *C. gasperettii* was recovered as the basal species (bootstrap values higher than 95%; Fig. S4). Within *C. cerastes* there were three different lineages: a northwestern African lineage (*C. c. cerastes*), a northeastern African lineage (*C. c. cerastes*) with two individuals from Chad and Egypt and a third lineage with the Arabian samples (*C. c. hoofieni*) sister to the northeastern African lineage. For *C. vipera*, the topology followed a geographical distribution of the four individuals sampled, with closer individuals clustering together. Conversely, in *C. gasperettii* there were two different lineages, splitting coastal and interior individuals from the Arabian Peninsula (Fig. S4). The time-calibrated Bayesian tree further corroborated the ML topology for the genus and provided support for the intraspecific relationships (Fig. 3B). The split between *C. gasperettii* and the ancestor of both *C. cerastes* and *C. vipera* was estimated around 7.12 ± 3.83 Mya and between *C. cerastes* and *C. vipera* around 4.22 ± 2.23 Mya (Fig. 3B). The genomic species tree inferred with SNAPP yielded the same new topology (Fig. S5), as well as the tree including *C. c. hoofieni* (Fig. S6) and divergence among species were found within the 95% interval of our time-calibrated Bayesian tree. Overall, intraspecific relationships were found to be consistent between mitochondrial and nuclear phylogenies (Fig. 3).

### Niche overlap analyses

The first three components accounted for more than 81% of the environmental variance in the two global PCAs, while this value was 60% in the regional-landcover PCA (Table S5). In the global-climate approach, *C. gasperettii* showed a much-reduced volume in comparison to the volume exhibited by the clade comprising *C. cerastes* and *C. vipera*, and also to the volumes recovered for each species. Conversely, in the global-landcover approach, *C. gasperettii* showed a similar volume to the volumes of the other two species, while it was more reduced when both species were considered as a single clade (Table S5). In the three approaches, *C. cerastes* and *C. vipera* showed rather similar volumes (Table S5).

Niche volume overlap was high in all cases (Fig. 4), with both K and OI indexes ranging from moderate to high (0.49 < K < 0.81; 0.70 < OI < 0.84; Table S5). Statistical tests were all significant (*p* < 0.05), with the exception of the test of K index for *C. cerastes* vs *C. vipera* in the global-landcover dataset (Table S5). Overall, niche overlap analyses show that the three species exhibit non-equivalent but highly overlapping niches.

**Fig. 4:**
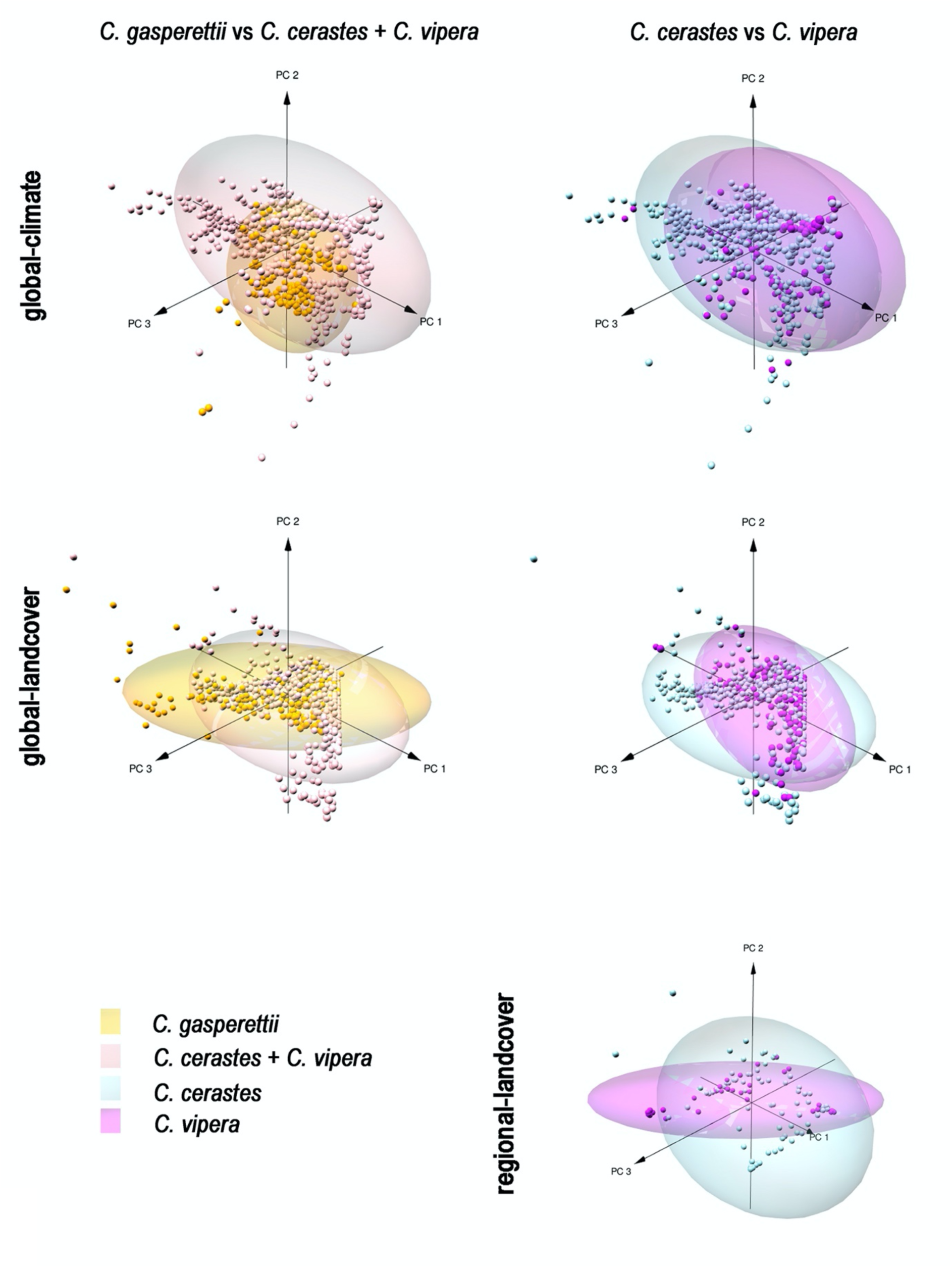
3D niche overlap plots. Each of the five plots represents the three first components of a principal component analyses (PCA) of the climatic/landcover variability for the occurrence of major clades found within *Cerastes*, approached at the global and regional scales. Ellipses represent the 0.95 confidence interval for the scores of each clade in the PCA.

### Introgression analyses

We used the filtered *SNP* dataset containing 32,265 SNPs that were present in at least 60% of the samples to study introgression between the different clades. Both D-statistics and f4-ratio analyses showed a significant excess of shared alleles between *C. vipera* and *C. c. cerastes* individuals (Table 1 and Fig. 5). Moreover, f-branch analyses supported D-statistics and f4-ratio and suggested an introgression of around 10% of the sampled sites between *C. vipera* and *C. c. cerastes* (Fig. 5). Ancestral introgression was not found between *C. gasperettii* and *C. c. hoofieni*, both occurring in Arabia (Table 1 and Fig. 5).

**Fig. 5:**
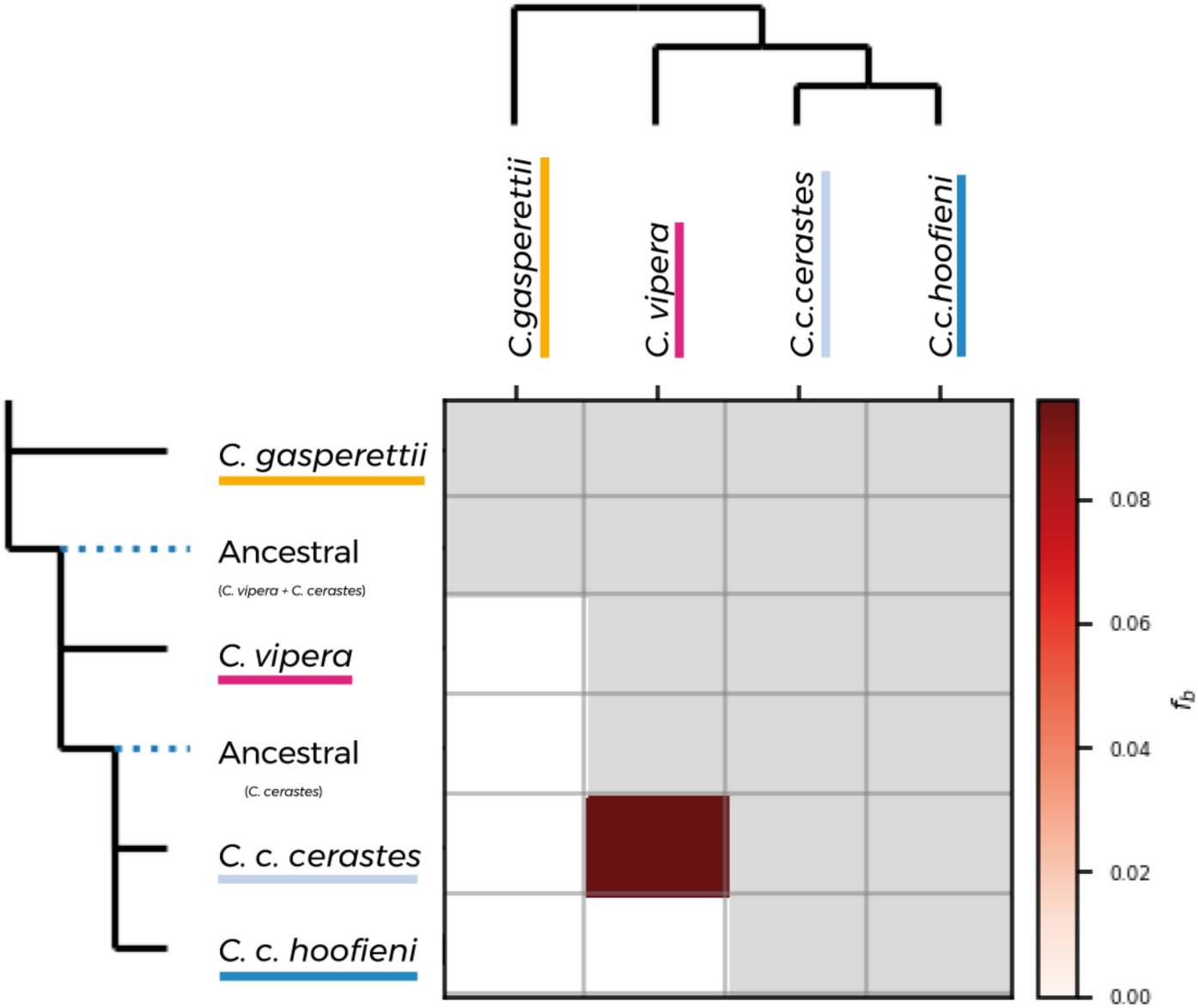
F-branch analysis indicating an ancestral introgression between the samples of *C. vipera* with *C. c. cerastes* over a total dataset of 32,254 SNPs, including *E. omanensis* as an outgroup.

**Table 1:** Introgression results for all the species and subspecies available in this study. Significant results are shown in bold. Abbreviations are as follows: Cch, *Cerastes cerastes hoofieni*; Ccc, *Cerastes cerastes cerastes*; Cg, *Cerastes gasperettii* and Cv, *Cerastes vipera*.

| P1 | P2 | P3 | D-statistics | Z-score | p-value | f4-ratio | BBAA | ABBA | BABA |
| --- | --- | --- | --- | --- | --- | --- | --- | --- | --- |
| Cch | Ccc | Cg | 0.03 | 0.57 | 0.28 | 0.004 | 631.28 | 60.44 | 56.08 |
| Cch | Ccc | Cv | 0.20 | **4.02** | **2.87e<sup>-5</sup>** | 0.096 | 331.97 | 131.93 | 87.66 |
| Cv | Ccc | Cg | 0.08 | 1.46 | 0.07 | 0.015 | 451.34 | 92.73 | 78.74 |
| Cch | Cv | Cg | 0.02 | 0.34 | 0.36 | 0.004 | 403.88 | 93.49 | 89.43 |

## Discussion

In this study, we used mitochondrial and genome-wide data to unveil the phylogenetic relationships of the genus *Cerastes*, a clade of medically important venomous snakes adapted to arid environments and mainly distributed across the Saharan and Arabian deserts. This group has been historically under- studied due to their wide distribution and harsh and remote environments it inhabits, which hampered a systematic field sampling. Our results unravel an unexpected phylogenomic relationship that differs from previous morphological and mitochondrial approaches, which suggested that *C. cerastes* and *C. gasperettii*, two morphologically similar species with shared habitat requirements, were sister taxa while, *C. vipera*, smaller and linked to soft sand habitats, was sister to them. Moreover, niche overlap analyses did not seem to support a potential scenario of sympatric speciation between *C. cerastes* and *C. vipera*. Ultimately, our study underscores the significance of employing genome-wide data in the examination of speciation processes and the unravelling of the complete evolutionary history of species. It also highlights the contrasting patterns observed between genomic data with both morphology and mitochondrial markers.

### Unexpected phylogenomic relationships within the genus *Cerastes*

All our phylogenomic analyses support a new phylogenomic relationship within the genus *Cerastes*. According to our findings*, C. cerastes* and *C. vipera* are sister taxa, while *C. gasperettii* is more distantly related (Fig. 3B and Fig. S4-S6). This contrasts with the previously accepted sister relationship between *C. cerastes* and *C. gasperettii*, with *C. vipera* considered as an external group relative to this clade (Alencar et al., 2016; Šmíd & Tolley, 2019; Fig. 3A). Population structure analyses also showed a similar clustering (Fig. 2). This was an unexpected result, since *C. cerastes* and *C. gasperettii* exhibit greater morphological similarities, both with horn and hornless populations, a similar body size, and overlapping habitat requirements, since both species are generalists (Werner et al., 1991; Brito et al., 2011). Conversely, *C. vipera* is smaller, strictly hornless and specialized in burrowing in sandy substrates, with specific adaptations such as the dorsal position of the eyes (Young & Morain, 2003; Sivan et al., 2013). This morphological-genomic contrasting pattern has also been reported for other species with complex evolutionary histories present in desert environments as foxes (Leite et al., 2015; Rocha et al. 2023).

Prior to our work, all phylogenetic knowledge on these desert-adapted vipers pointed towards *C. cerastes* and *C. gasperettii* being sister taxa (Pyron et al. 2013; Alencar et al., 2016; Šmíd & Tolley, 2019 and Fig. 3A). However, these studies predominantly relied on a variety of mitochondrial markers, since nuclear data has not been traditionally considered within the Viperinae subfamily due to its low variability (e.g., in Eurasian vipers, Freitas et al., 2020). One plausible hypothesis to consider is that the introgression observed between *C. c. cerastes* and *C. vipera* could have altered the phylogenetic position of *C. vipera*, potentially bringing it closer to *C. c. cerastes*. However, if that was the case, an incorporation of *C. c. hoofieni* into a phylogenetic framework should result in *C. cerastes* as paraphyletic (with *C. c. cerastes* and *C. vipera* clustering together and *C. c. hoofieni* closer to *C. gasperettii*), as we did not report introgression between *C. vipera* and the Arabian subspecies. Nonetheless, this was not the case as coancestry analyses would show more shared ancestry between *C. cerastes* and *C. gasperettii* (Fig. 2C) and the phylogenomic trees incorporating *C. c. hoofieni* also showed a common evolutionary history within *C. cerastes*, as both subspecies clustered together (Fig. 3, Fig. S4 and S6). This points towards incomplete lineage sorting of the mitochondria as the main factor contributing to the observed mitonuclear discordance described here. As it has been reported elsewhere, the use of nuclear (and especially genome-wide data) has been a key factor in unraveling the phylogenomic relationships among species (e.g., Mao et al., 2019). Mitochondrial data might not represent the complete evolutionary history of a species, as it only shows the evolutionary history of the maternally inherited mitochondria. Therefore, it is recommended to combine NGS techniques together with nuclear data to effectively address these questions.

### Speciation drivers in *Cerastes*

Vicariance, the geographic separation of connected populations eventually leading to different species due to the emergence of a physical barrier, is considered one of the key mechanisms driving speciation in low-dispersal reptiles from arid environments (Brito et al., 2014, Tejero-Cicuéndez et al., 2022b). This type of speciation has been repeatedly reported for species from the Arabian Peninsula and North Africa, as these areas present several geographical (e.g., environmental or physiographical) barriers that have had a determinant impact on the distribution and divergence of their reptile fauna (Brito et al., 2014; Metallinou et al., 2015; Garcia-Porta et al. 2017; Gonçalves et al., 2018a; 2018b; Liz et al., 2021; 2023; Tejero-Cicuéndez et al., 2022b; Burriel-Carranza et al., 2023b). Among them, deserts are an interesting geographical feature, as they can act both as a geographical barrier or a corridor, depending on the species’ ecological preference (Brito et al., 2014). On the one hand, deserts can serve as effective geographical barriers for several reptile species with rather mesic requirements, preventing gene flow between the populations on either side and leading to different evolutionary lineages (Rato et al., 2007; Martínez-Freiría et al., 2017; Gonçalves et al., 2018a; 2018b; Liz et al. 2021). On the other hand, deserts have facilitated the dispersal of arid-adapted taxa, expanding the range of xeric species during periods of increasing aridity (Amer & Kumazawa, 2005; Brito et al., 2014; Velo-Antón et al., 2018; Liz et al. 2021; 2023). In fact, during the mid-end of the Miocene, several arid-adapted genera such as *Hemidactylus*, *Acanthodactylus*, *Echis*, *Pseudotrapelus* or *Uromastyx* dispersed from Arabia to Africa, confirming the establishment of a dispersal corridor connecting Africa and Eurasia through the Arabian Peninsula (Tejero-Cicuéndez et al., 2022b). These dispersal events match with a progressive aridification of North Africa and Arabia during the mid-late Miocene, with an hyperaridity period in Arabia, together with the more recent phases of the formation of the Sahara Desert, which began around 7 Mya in northeastern Africa (Schuster et al., 2006; Böhme et al., 2021; Crocker et al., 2022).

Assuming an Arabian origin for the genus *Cerastes*, the rise of aridity and the formation of the Sahara Desert during the mid-late Miocene could have facilitated the colonization of North Africa by the arid-adapted ancestor of *C. cerastes* and *C. vipera*, a phenomenon previously observed in other species of reptiles (Amer & Kumazawa, 2005; Metallinou et al., 2012; Liz et al. 2021). In fact, this period coincides with the split of *C. gasperettii* and the ancestor of *C. cerastes* and *C. vipera*. Although Africa and Arabia started splitting around 30-25 Mya (Bohannon 1986; Ghebreab 1998; Bosworth et al., 2005) these two geographical entities were still in contact through the Sinai Peninsula (Bosworth et al., 2005; Tejero-Cicuéndez et al., 2022b). The subsequent formation of the Red Sea and the Gulf of Aden likely isolated *C. gasperettii* in Arabia, as these geographical barriers have caused strong vicariance speciation in other reptile species (Metallinou et al., 2012; Pook et al. 2009; Tejero-Cicuéndez et al., 2022b).

The dispersal of *Cerastes* from Arabia to Africa is relatively recent when compared to other taxa such as *Stenodactylus* or *Echis*, whose estimated colonization events dated around 21.8 and 19.4 Mya, respectively (Metallinou et al., 2012; Pook et al., 2009). However, there have been other dispersal events of species between Africa and Arabia that suggest that dispersion have also occurred at different times (e.g., 9.8 and 5.9 Mya for *Hemidactylus*, 8 Mya for *Echis pyramidum*, 4 Mya for *Bitis arietans* or 1.75 Mya for *Naja haje*; Pook et al., 2009; Šmíd et al., 2013). After the colonization of Africa, the dry-wet cycles that the Sahara experienced during the Plio-Pleistocene (Le Houérou, 1997; Brito et al., 2014) could have caused an isolation of the widespread *Cerastes* ancestor and a subsequent speciation into *C. cerastes* and *C. vipera*, around 4 Mya (Fig. 3B). Similar processes have been reported for other arid-adapted species included in the genera *Agama* (Gonçalves et al., 2012) or *Scincus* (Carranza et al., 2008; Šmíd et al., 2021b) and it is in line with our niche overlap analyses, as we did not find strong ecological differences between these two species (Fig. 4). This suggests that both species overlap in their habitat use and putatively contradicts a possible scenario of sympatric speciation. This is surprising, as *C. vipera* has been known to exhibit certain morphological adaptations, such as the dorsal position of the eyes that allows them to burrow in sandy substrates while ambushing potential preys (Young & Morain, 2003; Sivan et al., 2013). Conversely, *C. cerastes* has always been observed in rocky-like deserts (see Brito et al., 2011). The generalist ecological niche reported here for *C. cerastes* may lead to overlapping habitat use between both species (Fig. 4) or indicate that both species exploit the same ecological niche differently, as the digging habits present in *C. vipera*. Nonetheless, these unexpected ecological findings could also be attributed to a general absence of sampling of these unexplored areas, as recent surveys have found *C. vipera* in zones with compact sand or *C. cerastes* in sandy areas without rocks (authors, *pers. observation*). An alternate interpretation might be that we did not manage to capture the grain of the habitat at the scale we describe it as rocky/hard and sandy/soft are often present side by side.

The confirmation of the presence of *C. cerastes* in Arabia using genetic and genomic data (i.e., *C. c. hoofieni*), with its dispersal occurring around 1.7 Mya (Fig. 3B), could potentially be explained by either northern or southern land bridges. Currently, *C. c. hoofieni* is exclusively found in the coastal Tihama Desert in southwestern Arabia. In the scenario of a northern land bridge, it would suggest the species’ disappearance from the northwestern Saudi Arabia coastline. Conversely, a southern land bridge would imply its extirpation from Eritrea and Djibouti, a hypothesis that appears more plausible due to competition with other viperid snakes such as members of the genus *Echis*. To completely unravel the colonization route of *C. c. hoofieni*, it is essential to conduct a more thorough sampling (including individuals from both Sinai and southern Sudan) as well as bioclimatic models encompassing both current and past periods to unveil potential corridors for the species dispersal (e.g., Gonçalves et al., 2018a; Velo-Antón et al., 2018).

### Ancient introgression in North Africa

We found evidence of ancestral introgression between *C. c. cerastes* and *C. vipera*, which aligns with the high niche overlap reported here (Fig. 4 and 5 and Table 1). This introgression is supported by the different ancestry observed between *C. c. cerastes* and *C. c. hoofieni* with regards to *C. vipera* and most likely is due to the allopatric distribution of *C. c. hoofieni* and *C. vipera.* FineRADstructure also supported this finding, as *C. vipera* individuals shared high coancestry with *C. c. cerastes* samples (Fig. 2C). This discovery opens up avenues for future research to investigate the introgressed DNA regions and how this introgression could have affected the populations of *C. c. cerastes* compared to *C. c. hoofieni*, as introgression has been suggested as a source of adaptive variation (Hedrick, 2013) and has been proven fundamental for the arid adaption of some species of mammals (Liu et al., 2021; Nanaei et al. 2023; Rocha et al. 2023). Undoubtedly, additional individuals of *C. vipera* from northeastern Africa should be sampled to verify whether this introgression is maintained along the whole species’ range.

### Insights into the systematics of *Cerastes cerastes*

This study provides novel genetic and genomic information for the subspecies C*. c. hoofieni*. Moreover, our genome-wide data does not recognize *C. c. mutila* as an independent genetic unit, as it did not exhibit any genetic differences compared to other individuals of *C. c. cerastes* (Fig. 2-3 and Fig. S4). This finding further refutes the previous discussion on the validity of *C. c. mutila* as a subspecies (Martínez del Mármol et al., 2019). Conversely, our relatively small dataset seems to support the recognition of *C. c. hoofieni* as a subspecies within *C. cerastes* (Fig. 2-3 and Fig. S4), with *C. c. cerastes* being paraphyletic. Nevertheless, in spite of the existence of morphological differences (Werner et al., 1991), our genomic data so far does not support an elevation to a species status of *C. c. hoofieni* (Fig. 2-3 and Fig. S3), but future studies should address that with a bigger sampling and using an integrative taxonomic approach which combines morphological and climatic data together with genetics (e.g., Martínez-Freiría et al., 2021). Interestingly, the western populations of *C. cerastes* show a different genetic and evolutionary history than the rest, prompting the need for a taxonomic revision within this genetic cluster. Additionally, the taxonomic status of *C. g. mendelssohni* should be addressed in future research, as we did not manage to obtain samples for this study, although Carné et al., (2020) did not find genetic differences between this and the nominotypical subspecies using mitochondrial markers.

## Conclusions

In this study, we have substantially increased our understanding on the evolutionary history of *Cerastes* from the North African and Arabian regions. We provide, for the first time, genome-wide information for four medically important venomous snakes from North Africa and Arabia (including genome-wide data for *Echis omanensis*), as well as an unexpected phylogenomic relationship within the genus *Cerastes*, a crucial step for understanding the speciation processes that have shaped its evolutionary history. Furthermore, we have identified distinct genetic clades within *C. cerastes*. This finding is of particular significance in the study of venomous snakes, given the widespread nature of venom variation across different taxonomic levels (Chippaux et al., 1991; Rao et al., 2022). Ultimately, our study emphasizes the importance of employing nuclear and specifically genome-wide data, together with ecological data, to comprehensively infer evolutionary relationships between and within species.

## Acknowledgements

This work was supported by grants PGC2018-098290-B-I00 (MCIU/AEI/FEDER, UE), Spain, PID2021-128901NB-I00 (MCIN/AEI/10.13039/501100011033 and by ERDF, A way of making Europe), Spain, and grant 2021-SGR-00751 from the Departament de Recerca i Universitats from the Generalitat de Catalunya, Spain to SC. GM-R is supported by an FPI grant from the Ministerio de Ciencia, Innovación y Universidades, Spain (PRE2019-088729), BB-C is supported by FPU grant from Ministerio de Ciencia, Innovación y Universidades, Spain (FPU18/04742), AT is supported by “la Caixa” doctoral fellowship program (LCF/BQ/DR20/11790007), HT-C is supported by a “Juan de la Cierva - Formación” postdoctoral fellowship (FJC2021-046832-I) funded by MCIN/AEI/10.13039/501100011033 and by the European Union NextGenerationEU/PRTR, GV-A was supported by the FCT (CEECIND/00937/2018) and recently by a Ramón y Cajal research grant (Ref. RYC-2019-026959-I/AEI/10.13039/501100011033), JŠ was supported by the Czech Science Foundation (GAČR) under grant number 22-12757S and by the Charles University Research Centre under grant number 204069 and FM-F and JCB are supported by FCT - Fundação para a Ciencia e Tecnologia de Portugal (DL57/2016/CP1440/CT0010, CEECINST/00014/2018/CP1512/CT0001, respectively) and

## Data availability

All new COI sequences can be accessed at NCBI with Accession number SUB13696935 and demultiplex ddRAD sequencing files can be found under BioProject ID PRJNA997916.

## References

Alencar, L. R., Quental, T. B., Grazziotin, F. G., Alfaro, M. L., Martins, M., Venzon, M., & Zaher, H. (2016). Diversification in vipers: Phylogenetic relationships, time of divergence and shifts in speciation rates. Molecular phylogenetics and evolution, 105, 50–62.

Alexander, D. H., Novembre, J., & Lange, K. (2009). Fast model-based estimation of ancestry in unrelated individuals. Genome Research, 19(9), 1655–1664. 10.1101/GR.094052.109

Amer, S. A., & Kumazawa, Y. (2005). Mitochondrial DNA sequences of the Afro-Arabian spiny-tailed lizards (genus Uromastyx; family Agamidae): phylogenetic analyses and evolution of gene arrangements. Biological Journal of the Linnean Society, 85(2), 247–260.

Anderson, T.M., vonHoldt, B.M., Candille, S.I., Musiani, M., Greco, C., Stahler, D.R., Smith, D.W., Padhukasahasram, B., Randi, E., Leonard, J.A. and Bustamante, C.D. (2009). Molecular and evolutionary history of melanism in North American gray wolves. Science, 323(5919), pp.1339–1343. 10.1126/science.1165448

Andrews, S. (2010). FastQC: a quality control tool for high throughput sequence data.

Barraclough, T. G., & Nee, S. (2001). Phylogenetics and speciation. Trends in Ecology & Evolution, 16(7), 391–399. 10.1016/S0169-5347(01)02161-9

Bicheron, P., Defourny, P., Brockmann, C., Schouten, L., Vancutsem, C., Huc, M., Bontemps, S., Leroy, M., Achard, F., Herold, M., Ranera, F., Arino, O., 2008. GLOBCOVER: Products Description and Validation Report. http://postel.Mediasfrance.org Medias-France and Postel.

Blonder, B., Lamanna, C., Violle, C., & Enquist, B. J. (2014). The n-dimensional hypervolume. Global Ecology and Biogeography, 23(5), 595–609.

Bohannon, R. W. (1986). Test-retest reliability of hand-held dynamometry during a single session of strength assessment. Physical therapy, 66(2), 206–209. 10.1093/ptj/66.2.206

Böhme, M., Spassov, N., Ebner, M., Geraads, D., Hristova, L., Kirscher, U., … & Winklhofer, M. (2017). Messinian age and savannah environment of the possible hominin Graecopithecus from Europe. PloS one, 12(5), e0177347.

Bosworth, W., Khalil, S. M., Ligi, M., Stockli, D. F., & McClay, K. R. (2020). Geology of Egypt: The Northern Red Sea. The geology of Egypt, 343-374. 10.1007/978-3-030-15265-9

Bouckaert, R. (2010). DensiTree: Making sense of sets of phylogenetic trees. Bioinformatics, 26(10), 1372–1373. 10.1093/bioinformatics/btq110

Bouckaert, R., Vaughan, T. G., Barido-Sottani, J., Duchêne, S., Fourment, M., Gavryushkina, A., Heled, J., Jones, G., Kühnert, D., De Maio, N., Matschiner, M., Mendes, F. K., Müller, N. F., Ogilvie, H. A., Du Plessis, L., Popinga, A., Rambaut, A., Rasmussen, D., Siveroni, I., … Drummond, A. J. (2019). BEAST 2.5: An advanced software platform for Bayesian evolutionary analysis. PloS Computational Biology, 15(4). 10.1371/journal.pcbi.1006650

Brito, J. C., Godinho, R., Martínez-Freiría, F., Pleguezuelos, J. M., Rebelo, H., Santos, X., Vale, C. G., Velo-Antón, G., Boratyński, Z., Carvalho, S. B., Ferreira, S., Gonçalves, D. V., Silva, T. L., Tarroso, P., Campos, J. C., Leite, J. V., Nogueira, J., Álvares, F., Sillero, N., … Carranza, S. (2014). Unravelling biodiversity, evolution and threats to conservation in the Sahara-Sahel: Biodiversity patterns in the Sahara-Sahel. Biological Reviews, 89(1), 215–231. 10.1111/brv.12049

Brito, J. C., Fahd, S., Geniez, P., Martínez-Freiría, F., Pleguezuelos, J. M., & Trape, J. F. (2011). Biogeography and conservation of viperids from North-West Africa: an application of ecological niche-based models and GIS. Journal of Arid Environments, 75(11), 1029–1037.

Brown, T. (2002). Genomes.

Bryant, D., Bouckaert, R., Felsenstein, J., Rosenberg, N. A., & Roychoudhury, A. (2012). Inferring species trees directly from biallelic genetic markers: Bypassing gene trees in a full coalescent analysis. Molecular Biology and Evolution, 29(8), 1917–1932. 10.1093/molbev/mss086

Burriel-Carranza B., Tarroso P., Els J., Gardner A., Soorae P., Mohammed A.A., Tubati S.R.K., Eltayeb M.M., Shah J.N., Tejero-Cicuéndez H., Simó-Riudalbas M., Pleguezuelos J.M., Fernández-Guiberteau D., Šmíd J., Carranza S. 2019. An integrative assessment of the diversity, phylogeny, distribution, and conservation of the terrestrial reptiles (Sauropsida, Squamata) of the United Arab Emirates. PLOS ONE. 14:e0216273. 10.1371/journal.pone.0216273

Burriel-Carranza, B., Estarellas, M., Riaño, G., Talavera, A., Tejero-Cicuéndez, H., Els, J., & Carranza, S. (2023a). Species boundaries to the limit: Integrating species delimitation methods is critical to avoid taxonomic inflation in the case of the Hajar banded ground gecko (*Trachydactylus hajarensis*). Molecular Phylogenetics and Evolution, 186, 107834. 10.1016/j.ympev.2023.107834

Burriel-Carranza, B., Tejero-Cicuéndez, H., Carné, A., Riaño, G., Talavera, A., Al Saadi, S., Els, J., Šmíd, J., Tamar, K., Tarroso, P., Carranza, S. (2023b). The origin of a mountain biota: hyper- aridity shaped reptile diversity in an Arabian biodiversity hotspot. bioRxiv 2023.04.07.536010. doi: 10.1101/2023.04.07.536010

Cahill, J. A., Stirling, I., Kistler, L., Salamzade, R., Ersmark, E., Fulton, T. L., Stiller, M., Green, R. E., & Shapiro, B. (2015). Genomic evidence of geographically widespread effect of gene flow from polar bears into brown bears. Molecular Ecology, 24(6), 1205–1217. 10.1111/mec.13038

Campos, J. C., & Brito, J. C. (2018). Mapping underrepresented land cover heterogeneity in arid regions: the Sahara-Sahel example. ISPRS Journal of Photogrammetry and Remote Sensing, 146, 211–220.

Carné, A., Fathinia, B., & Rastegar-Pouyani, E. (2020). Molecular phylogeny of the Arabian Horned Viper, *Cerastes gasperettii* (Serpentes: Viperidae) in the Middle East. Zoology in the Middle East, 66(1), 13–20. 10.1080/09397140.2020.1711622

Carranza, S., Arnold, E. N., Geniez, P., Roca, J., & Mateo, J. A. (2008). Radiation, multiple dispersal and parallelism in the skinks, Chalcides and Sphenops (Squamata: Scincidae), with comments on Scincus and Scincopus and the age of the Sahara Desert. Molecular phylogenetics and evolution, 46(3), 1071–1094. 10.1016/j.ympev.2007.11.018

Carranza, S., Xipell, M., Tarroso, P., Gardner, A., Arnold, E. N., Robinson, M. D., Simó-Riudalbas, M., Vasconcelos, R., de Pous, P., Amat, F., Šmíd, J., Sindaco, R., Metallinou, M., Els, J., Pleguezuelos, J. M., Machado, L., Donaire, D., Martínez, G., Garcia-Porta, J., … Al Akhzami, S. N. (2018). Diversity, distribution and conservation of the terrestrial reptiles of Oman (Sauropsida, Squamata). PLOS ONE, 13(2), e0190389. 10.1371/journal.pone.0190389

Chang, C. C., Chow, C. C., Tellier, L. C. A. M., Vattikuti, S., Purcell, S. M., & Lee, J. J. (2015). Second-generation PLINK: Rising to the challenge of larger and richer datasets. GigaScience, 4(1). 10.1186/s13742-015-0047-8

Chippaux, J. P., Williams, V., & White, J. (1991). Snake venom variability: methods of study, results and interpretation. Toxicon, 29(11), 1279–1303. 10.1016/0041-0101(91)90116

Crocker, A. J., Naafs, B. D. A., Westerhold, T., James, R. H., Cooper, M. J., Röhl, U., Pancost, R. D., Xuan, C., Osborne, C. P., Beerling, D. J., & Wilson, P. A. (2022). Astronomically controlled aridity in the Sahara since at least 11 million years ago. Nature Geoscience, 15(8), Article 8. 10.1038/s41561-022-00990-7

Dinis, M., Merabet, K., Martínez-Freiría, F., Steinfartz, S., Vences, M., Burgon, J. D., … & Velo-Antón, G. (2019). Allopatric diversification and evolutionary melting pot in a North African Palearctic relict: the biogeographic history of *Salamandra algira*. Molecular Phylogenetics and Evolution, 130, 81–91.

Eaton, D. A. R., & Overcast, I. (2020). Ipyrad: Interactive assembly and analysis of RADseq datasets. Bioinformatics, 36(8), 2592–2594. 10.1093/bioinformatics/btz966

Freitas, I., Ursenbacher, S., Mebert, K., Zinenko, O., Schweiger, S., Wüster, W., … & Martínez-Freiría, F. (2020). Evaluating taxonomic inflation: towards evidence-based species delimitation in Eurasian vipers (Serpentes: Viperinae). Amphibia-Reptilia, 41(3), 285–311.

García-Cardenete, L., Flores-Stols, M. V., & Yubero, S. (2017). New cases of syntopy between viperid snakes (Viperidae) in the Atlantic Sahara. Go-South Bulletin, 14, 139–141.

Garcia-Porta, J., Simó-Riudalbas, M., Robinson, M., & Carranza, S. (2017). Diversification in arid mountains: Biogeography and cryptic diversity of *Pristurus rupestris rupestris* in Arabia. Journal of Biogeography, 44(8), 1694–1704. 10.1111/jbi.12929

Ghebreab, W. (1998). Tectonics of the Red Sea region reassessed. Earth-Science Reviews, 45(1-2), 1–44. 10.1016/S0012-8252(98)00036-1

Gonçalves, D. V., Brito, J. C., Crochet, P. A., Geniez, P., Padial, J. M., & Harris, D. J. (2012). Phylogeny of North African *Agama* lizards (Reptilia: Agamidae) and the role of the Sahara desert in vertebrate speciation. Molecular phylogenetics and evolution, 64(3), 582–591. 10.1016/j.ympev.2012.05.007

Gonçalves, D. V., Martínez-Freiría, F., Crochet, P. A., Geniez, P., Carranza, S., & Brito, J. C. (2018a). The role of climatic cycles and trans-Saharan migration corridors in species diversification: biogeography of *Psammophis schokari* group in North Africa. Molecular Phylogenetics and Evolution, 118, 64–74.

Gonçalves, D. V., Pereira, P., Velo-Antón, G., Harris, D. J., Carranza, S., & Brito, J. C. (2018b). Assessing the role of aridity-induced vicariance and ecological divergence in species diversification in North-West Africa using Agama lizards. Biological Journal of the Linnean Society, 124(3), 363–380. 10.1093/biolinnean/bly055

Gosselin, T., Lamothe, M., Devloo-Delva, F., & Grewe, P. (2017). Radiator: RADseq data exploration, manipulation and visualization using R. R package version 0.0. 5. Retrieved from https://github.Com/25heirrygosselin/radiator.

Green, R. E., Krause, J., Briggs, A. W., Maricic, T., Stenzel, U., Kircher, M., Patterson, N., Li, H., Zhai, W., Fritz, M. H.-Y., Hansen, N. F., Durand, E. Y., Malaspinas, A.-S., Jensen, J. D., Marques-Bonet, T., Alkan, C., Prüfer, K., Meyer, M., Burbano, H. A., … Pääbo, S. (2010). A Draft Sequence of the Neandertal Genome. Science, 328(5979), 710–722. 10.1126/science.1188021

Guo, P., Liu, Q., Zhu, F., Zhong, G. H., Che, J., Wang, P., Xie, Y. L., Murphy, R. W., & Malhotra, A. (2019). Multilocus phylogeography of the brown-spotted pitviper *Protobothrops mucrosquamatus* (Reptilia: Serpentes: Viperidae) sheds a new light on the diversification pattern in Asia. Molecular Phylogenetics and Evolution, 133, 82–91. 10.1016/j.ympev.2018.12.028

Harrison, R. G., & Larson, E. L. (2014). Hybridization, Introgression, and the Nature of Species Boundaries. Journal of Heredity, 105(S1), 795–809. 10.1093/jhered/esu033

Hedrick, Philip W. (2013). Adaptive introgression in animals: examples and comparison to new mutation and standing variation as sources of adaptive variation. Molecular ecology, 22(18) 4606–4618. 10.1111/mec.12415

Hinojosa, J. C., Koubínová, D., Szenteczki, M. A., Pitteloud, C., Dincă, V., Alvarez, N., & Vila, R. (2019). A mirage of cryptic species: Genomics uncover striking mitonuclear discordance in the butterfly *Thymelicus sylvestris*. Molecular Ecology, 28(17), 3857–3868. 10.1111/mec.15153

Katoh, K., & Standley, D. M. (2013). MAFFT Multiple Sequence Alignment Software Version 7: Improvements in Performance and Usability. Molecular Biology and Evolution, 30(4), 772– 780. 10.1093/molbev/mst010

Knaus, B. J., & Grünwald, N. J. (2017). Vcfr: A package to manipulate and visualize variant call format data in R. Molecular Ecology Resources, 17(1), 44–53. 10.1111/1755-0998.12549

Kozlov, A. M., Darriba, D., Flouri, T., Morel, B., & Stamatakis, A. (2019). RaxML-NG: a fast, scalable and user-friendly tool for maximum likelihood phylogenetic inference. Bioinformatics, 35(21), 4453–4455. 10.1093/BIOINFORMATICS/BTZ305

Leite, J. V., Álvares, F., Velo-Antón, G., Brito, J. C., & Godinho, R. (2015). Differentiation of North African foxes and population genetic dynamics in the desert—insights into the evolutionary history of two sister taxa, Vulpes rueppellii and Vulpes vulpes. Organisms Diversity & Evolution, 15, 731–745. 10.1007/s13127-015-0232-8

Leonard, J. A., Echegaray, J., Rand, E., & Vilà, C. (2013). Impact of hybridization with domestic dogs on the conservation of wild canids. In M. E. Gompper (Ed.), Free-Ranging Dogs and Wildlife Conservation (pp. 170–184). Oxford University Press. 10.1093/acprof:osobl/9780199663217.003.0007

Le Houérou, H. N. (1997). Climate, flora and fauna changes in the Sahara over the past 500 million years. Journal of Arid Environments, 37(4), 619–647. 10.1006/jare.1997.0315

Liu, X., Li, Z., Yan, Y., Li, Y., Wu, H., Pei, J., Yan, P., Yang, R., Gui, X., & Lan, X. (2021). Selection and introgression facilitated the adaptation of Chinese native endangered cattle in extreme environments. Evolutionary Applications, 14(3), 860–873. 10.1111/eva.13168

Liz, A. V., Rödder, D., Gonçalves, D. V., Velo-Antón, G., Fonseca, M. M., Geniez, P., Crochet, P.- A.& Brito, J. C. (2021). The role of Sahara highlands in the diversification and desert colonization of the Bosc’s fringe-toed lizard. Journal of Biogeography, 48(11), 2891–2906. 10.1111/jbi.14250

Liz, A. V., Rödder, D., Gonçalves, D. V., Velo-Antón, G., Tarroso, P., Geniez, P., … & Brito, J. C. (2023). Overlooked species diversity in the hyper-arid Sahara Desert unveiled by dryland-adapted lizards. Journal of Biogeography, 50(1), 101–115. 10.1111/jbi.14510

Lucchini, N., Kaliontzopoulou, A., Lourdais, O., & Martínez-Freiría, F. (2023). Climatic adaptation explains responses to Pleistocene oscillations and diversification in European vipers. Journal of Biogeography.

Malinsky, M., Matschiner, M., & Svardal, H. (2021). Dsuite - Fast *D* -statistics and related admixture evidence from VCF files. Molecular Ecology Resources, 21(2), 584–595. 10.1111/1755-0998.13265

Malinsky, M., Trucchi, E., Lawson, D. J., & Falush, D. (2018). RADpainter and fineRADstructure: Population Inference from RADseq Data. Molecular Biology and Evolution, 35(5), 1284–1290. 10.1093/molbev/msy023

Martínez del Mármol, G., Harris, D.J., Geniez, P., de Pous, P., Salvi, D. (2019). Amphibians and Reptiles of Morocco.

Martínez-Freiría, F., Velo-Antón, G., & Brito, J. C. (2015). Trapped by climate: interglacial refuge and recent population expansion in the endemic Iberian adder Vipera seoanei. Diversity and Distributions, 21(3), 331–344. 10.1111/ddi.12265

Martínez-Freiría, F., Crochet, P. A., Fahd, S., Geniez, P., Brito, J. C., & Velo-Antón, G. (2017). Integrative phylogeographical and ecological analysis reveals multiple Pleistocene refugia for Mediterranean Daboia vipers in north-west Africa. Biological Journal of the Linnean Society, 122(2), 366–384. 10.1093/biolinnean/blx038

Martínez-Freiría, F., Freitas, I., Zuffi, M. A., Golay, P., Ursenbacher, S., & Velo-Antón, G. (2020). Climatic refugia boosted allopatric diversification in western Mediterranean vipers. Journal of Biogeography, 47(8), 1698–1713. 10.1111/jbi.13861

Martínez-Freiría, F., Freitas, I., Velo-Antón, G., Lucchini, N., Fahd, S., Larbes, S., Pleguezuelos, J.M., Santos, X. and Brito, J. C. (2021). (2021). Integrative taxonomy reveals two species and intraspecific differentiation in the Vipera latastei–monticola complex. Journal of Zoological Systematics and Evolutionary Research 59(8): 2278–2306. 10.1111/jzs.12534

Mao, X., Tsagkogeorga, G., Thong, V. D., & Rossiter, S. J. (2019). Resolving evolutionary relationships among six closely related taxa of the horseshoe bats (*Rhinolophus*) with targeted resequencing data. Molecular Phylogenetics and Evolution, 139, 106551. 10.1016/j.ympev.2019.106551

Meirmans, P. G. (2020). Genodive version 3.0: Easy-to-use software for the analysis of genetic data of diploids and polyploids. Molecular Ecology Resources, 20(4), 1126–1131. 10.1111/1755-0998.13145

Metallinou, M., Arnold, E. N., Crochet, P.-A., Geniez, P., Brito, J. C., Lymberakis, P., Baha El Din, S., Sindaco, R., Robinson, M., & Carranza, S. (2012). Conquering the Sahara and Arabian deserts: Systematics and biogeography of Stenodactylus geckos (Reptilia: Gekkonidae). BMC Evolutionary Biology, 12(1), 258. 10.1186/1471-2148-12-258

Metallinou, M., Červenka, J., Crochet, P. A., Kratochvíl, L., Wilms, T., Geniez, P., Shobrak, M. Y., Brito, J. C., & Carranza, S. (2015). Species on the rocks: Systematics and biogeography of the rock-dwelling Ptyodactylus geckos (Squamata: Phyllodactylidae) in North Africa and Arabia. Molecular Phylogenetics and Evolution, 85, 208–220. 10.1016/j.ympev.2015.02.010

Mochales-Riaño, G., Fontsere, C., de Manuel, M., Talavera, A., Burriel-Carranza, B., Tejero-Cicuéndez, H., AlGethami, R.H.M., Shobrak, M., Marques-Bonet, T. & Carranza, S. (2023). Genomics reveals introgression and purging of deleterious mutations in the Arabian leopard (Panthera pardus nimr). Iscience, 26(9). 10.1016/j.isci.2023.107481

Moutinho, A. F., Serén, N., Paupério, J., Silva, T. L., Martínez-Freiría, F., Sotelo, G., … & Boratyński, Z. (2020). Evolutionary history of two cryptic species of northern African jerboas. BMC Evolutionary Biology, 20, 1–16.10.1186/s12862-020-1592-z

Nagy, Z. T., Sonet, G., Glaw, F., & Vences, M. (2012). First Large-Scale DNA Barcoding Assessment of Reptiles in the Biodiversity Hotspot of Madagascar, Based on Newly Designed COI Primers. PloS ONE, 7(3), e34506. 10.1371/journal.pone.0034506

Nanaei, A.H., Cai, Y., Alshawi, A., Wen, J., Hussain, T., W.-W., Xu, N.Y., Essa, A., Lenstra, J.A., Wang, X., & Jiang, Y. (2023). Genomic analysis of indigenous goats in Southwest Asia reveals evidence of ancient adaptive introgression related to desert climate. Zoological Research, 44(1), 20–29. 10.24272/j.issn.2095-8137.2022.242

O’Leary, S. J., Puritz, J. B., Willis, S. C., Hollenbeck, C. M., & Portnoy, D. S. (2018). These aren’t the loci you’e looking for: Principles of effective SNP filtering for molecular ecologists. Molecular Ecology, 27(16), 3193–3206. 10.1111/MEC.14792

Ottenburghs, J., Megens, H.-J., Kraus, R. H. S., van Hooft, P., van Wieren, S. E., Crooijmans, R. P. M. A., Ydenberg, R. C., Groenen, M. A. M., & Prins, H. H. T. (2017). A history of hybrids? Genomic patterns of introgression in the True Geese. BMC Evolutionary Biology, 17(1), 201. 10.1186/s12862-017-1048-2

Peterson, B. K., Weber, J. N., Kay, E. H., Fisher, H. S., & Hoekstra, H. E. (2012). Double Digest RADseq: An Inexpensive Method for De Novo SNP Discovery and Genotyping in Model and Non-Model Species. PLOS ONE, 7(5), e37135. 10.1371/journal.pone.0037135

Phelps, T. (2010). Old world vipers: a natural history of the Azemiopinae and Viperinae. Edition Chimaira.

Pook, C. E., Joger, U., Stümpel, N., & Wüster, W. (2009). When continents collide: phylogeny, historical biogeography and systematics of the medically important viper genus Echis (Squamata: Serpentes: Viperidae). Molecular Phylogenetics and Evolution, 53(3), 792–807. 10.1016/j.ympev.2009.08.002

Pyron, R. A., Burbrink, F. T., & Wiens, J. J. (2013). A phylogeny and revised classification of Squamata, including 4161 species of lizards and snakes. BMC evolutionary biology, 13, 1–54.

R Core Team. (2021). R: A Language and Environment for Statistical Computing. https://www.R-project.org/

Rambaut, A., Drummond, A. J., Xie, D., Baele, G., & Suchard, M. A. (2018). Posterior Summarization in Bayesian Phylogenetics Using Tracer 1.7. Systematic Biology, 67(5), 901– 904. 10.1093/sysbio/syy032

Rao, W. Q., Kalogeropoulos, K., Allentoft, M. E., Gopalakrishnan, S., Zhao, W. N., Workman, C. T., … & Laustsen, A. H. (2022). The rise of genomics in snake venom research: recent advances and future perspectives. GigaScience, 11. 10.1093/gigascience/giac024

Rato, C., Brito, K.C., Carretero, M.A., Larbes, S., Shacham, B., Harris, D.J. (2007). Phylogeography and genetic diversity of *Psammophis schokari* (Serpentes) in North Africa based on mitochondrial DNA sequences. African Zoology, 42(1), 112–117 10.1080/15627020.2007.11407383

Rocha, J., Silva, P., Santos, N., Nakamura, M., Afonso, S., Qninba, A., Boratynksku, Z., Sudmant, P.H., Brito, J.C., Nielsen, R. & Godinho, R. (2023). North African fox genomes show signatures of repeated introgression and adaptation to life in deserts. Nature Ecology & Evolution, 7(8). 10.1038/s41559-023-02094-w

Roll, U., Feldman, A., Novosolov, M., Allison, A., Bauer, A. M., Bernard, R., Böhm, M., Castro-Herrera, F., Chirio, L., Collen, B., Colli, G. R., Dabool, L., Das, I., Doan, T. M., Grismer, L. L., Hoogmoed, M., Itescu, Y., Kraus, F., LeBreton, M., … Meiri, S. (2017). The global distribution of tetrapods reveals a need for targeted reptile conservation. Nature Ecology & Evolution, 1(11), 1677–1682. 10.1038/s41559-017-0332-2

Schoch, C. L., Sung, G.-H., López-Giráldez, F., Townsend, J. P., Miadlikowska, J., Hofstetter, V., Robbertse, B., Matheny, P. B., Kauff, F., Wang, Z., Gueidan, C., Andrie, R. M., Trippe, K., Ciufetti, L. M., Wynns, A., Fraker, E., Hodkinson, B. P., Bonito, G., Groenewald, J. Z., … Spatafora, J. W. (2009). The Ascomycota Tree of Life: A Phylum-wide Phylogeny Clarifies the Origin and Evolution of Fundamental Reproductive and Ecological Traits. Systematic Biology, 58(2), 224–239. 10.1093/sysbio/syp020

Schuster, M., Duringer, P., Ghienne, J. F., Vignaud, P., Mackaye, H. T., Likius, A., & Brunet, M. (2006). The age of the Sahara desert. Science, 311(5762), 821–821. 10.1126/science.1120161

Seehausen, O. (2004). Hybridization and adaptive radiation. Trends in Ecology & Evolution, 19(4), 198–207. 10.1016/j.tree.2004.01.003

Simó-Riudalbas, M., Tarroso, P., Papenfuss, T., Al-Sariri, T., & Carranza, S. (2018). Systematics, biogeography and evolution of *Asaccus gallagheri* (Squamata, Phyllodactylidae) with the description of a new endemic species from Oman. Systematics and Biodiversity, 16(4), 323–339.

Sindaco et al. 2014. The Reptiles of the Western Palearctic, Volume 2: Annotated Checklist and Distributional Atlas of the Snakes of Europe, North Africa, Middle East and Central Asia

Sivan, J., Kam, M., Hadad, S., Degen, A. A., Rozenboim, I., & Rosenstrauch, A. (2013). Temporal activity and dietary selection in two coexisting desert snakes, the Saharan sand viper (Cerastes vipera) and the crowned leafnose (Lytorhynchus diadema). Zoology, 116(2), 113–117. 10.1016/j.zool.2012.09.002

Šmíd, J., Carranza, S., Kratochvíl, L., Gvoždík, V., Nasher, A. K., & Moravec, J. (2013). Out of Arabia: A Complex Biogeographic History of Multiple Vicariance and Dispersal Events in the Gecko Genus Hemidactylus (Reptilia: Gekkonidae). PLoS ONE, 8(5), e64018. 10.1371/journal.pone.0064018

Šmíd, J., & Tolley, K. A. (2019). Calibrating the tree of vipers under the fossilized birth-death model. Scientific Reports, 9(1), 5510. 10.1038/s41598-019-41290-2

Šmíd, J., Sindaco, R., Shobrak, M., Busais, S., Tamar, K., Aghová, T., Simó-Riudalbas, M., Tarroso, P., Geniez, P., Crochet, P-A., Els, J., Burriel-Carranza, B., Tejero-Cicuéndez, H. & Carranza, S. (2021a). Diversity patterns and evolutionary history of Arabian squamates. Journal of Biogeography, 48(5), 1183–1199. 10.1111/jbi.14070

Šmíd, J., Uvizl, M., Shobrak, M., Salim, A. F. A., AlGethami, R. H. M., Algethami, A. R., Alanazi, A. S. K., Alsubaie, S. D., Busais, S., & Carranza, S. (2021b). Swimming through the sands of the Sahara and Arabian deserts: Phylogeny of sandfish skinks (Scincidae, Scincus) reveals a recent and rapid diversification. Molecular Phylogenetics and Evolution, 155, 107012. 10.1016/j.ympec.2020.107012

Song, Ying, Stefan Endepols, Nicole Klemann, Dania Richter, Franz-Rainer Matuschka, Ching-Hua Shih, Michael W. Nachman, and Michael H. Kohn. (2011). Adaptive introgression of anticoagulant rodent poison resistance by hybridization between old world mice. Current Biology, 21(15): 1296–1301. 10.1016/j.cub.2011.06.043

Stange, M., Sánchez-Villagra, M. R., Salzburger, W., & Matschiner, M. (2018). Bayesian divergence-time estimation with genome-wide single-nucleotide polymorphism data of sea catfishes (Ariidae) supports Miocene closure of the Panamanian Isthmus. Systematic Biology, 67(4), 681–699. 10.1093/sysbio/syy006

Tejero-Cicuéndez, H., Tarroso, P., Carranza, S., & Rabosky, D. L. (2022a). Desert lizard diversity worldwide: Effects of environment, time, and evolutionary rate. Global Ecology and Biogeography, 31(4), 776–790. 10.1111/geb.13470

Tejero-Cicuéndez, H., Patton, A. H., Caetano, D. S., Šmíd, J., Harmon, L. J., & Carranza, S. (2022b). Reconstructing Squamate Biogeography in Afro-Arabia Reveals the Influence of a Complex and Dynamic Geologic Past. Systematic Biology, 71(2), 261–272. 10.1093/sysbio/syab025

Thanou, E., Jablonski, D., & Kornilios, P. (2023). Genome-wide single nucleotide polymorphisms reveal recurrent waves of speciation in niche-pockets, in Europe’s most venomous snake. Molecular Ecology, 32(13), 3624–3640. 10.1111/mec.16944

Trape, J.-F. (2023). Guide des serpents d’Afrique occidentale, centrale et d’Afrique du Nord. IRD Editions, Marseille, 896 pp

Toews, D. P. L. & Brelsford, A. (2012). The biogeography of mitochondrial and nuclear discordance in animals. Molecular Ecology, 21(16), 3907–3930. 10.1111/j.1365-294X.2012.05664.x

Uetz, P., Freed, P, Aguilar, R., Reyes, F. & Hošek, J. (eds.) (2023) The Reptile Database, http://www.reptile-database.org, accessed [19/09/2023].

Velo-Antón, G., Martínez-Freiría, F. Pereira, P. Crochet, P.-A & Brito, J.C. (2018). Living on the edge:, Ecological and genetic connectivity of the spiny-footed lizard, *Acanthodactylus aureus*, confirms the Atlantic Sahara desert as a biogeographic corridor and centre of lineage diversification. Journal of Biogeography, 45(5), 1031–1042. 10.1111/jbi.13176

Velo-Antón, Guillermo, Margarida Henrique, André Vicente Liz, Fernando Martínez-Freiría, Juan Manuel Pleguezuelos, Philippe Geniez, Pierre-André Crochet, and José Carlos Brito. (2022). DNA barcode reference library for the West Sahara-Sahel reptiles. Scientific Data, 9(1): 459. 10.1038/s41597-022-01582-1

Wagner, P., & Wilms, T. (2010). A crowned devil: New species of *Cerastes* Laurenti, 1768 (Ophidia, Viperidae) from Tunesia, with two nomenclatural comments. Bonn Zoological Bulletin, 57, 297–306.

Warren, D. L., Glor, R. E., & Turelli, M. (2010). ENMTools: a toolbox for comparative studies of environmental niche models. Ecography, 33(3), 607–611.

Werner, Y., Le Verdier, A., Rosenman, D., & Sivan, N. (1991). Systematics and zoogeography of *Cerastes* (Ophidia: Viperidae) in the Levant: 1, Distinguishing Arabian from African “*Cerastes cerastes*.” The Snake, 23, 90–100.

Werner, Y., Sivan, N., Kushnir, V., & Motro, U. (1999). A statistical approach to variation in *Cerastes* (Ophidia: Viperidae), with the description of two endemic subspecies. Kaupia: Darmstädter Beiträge Zur Naturgeschichte, 8, 83–97.

Wickham, H. (2016). Ggplot2: Elegant Graphics for Data Analysis. Springer-Verlag New York. https://ggplot2.tidyverse.org

Young, B. A. & Morain, M. (2003). Vertical Burrowing in the Saharan Sand Vipers (*Cerastes*). Copeia, 2003(1), 131–137.

Zaidi, A. A., & Makova, K. D. (2019). Investigating mitonuclear interactions in human admixed populations. Nature Ecology & Evolution, 3(2), Article 2. 10.1038/s41559-018-0766-1

Zhang, Z., Jin, Y., Gao, Y., Zhang, Y., Ying, Q., Shen, C., Lu, J., Zhan, X., Wang, H., & Feng, S. (2022). The Complete Chloroplast Genomes of Two *Physalis* Species, *Physalis macrophysa* and *P. ixocarpa*: Comparative Genomics, Evolutionary Dynamics and Phylogenetic Relationships. Agronomy, 13(1), 135. 10.3390/agronomy13010135

