## Supplementary information for "Hidden in the sand: phylogenomics unravel an unexpected evolutionary history on the desert-adapted vipers of the genus *Cerastes*"

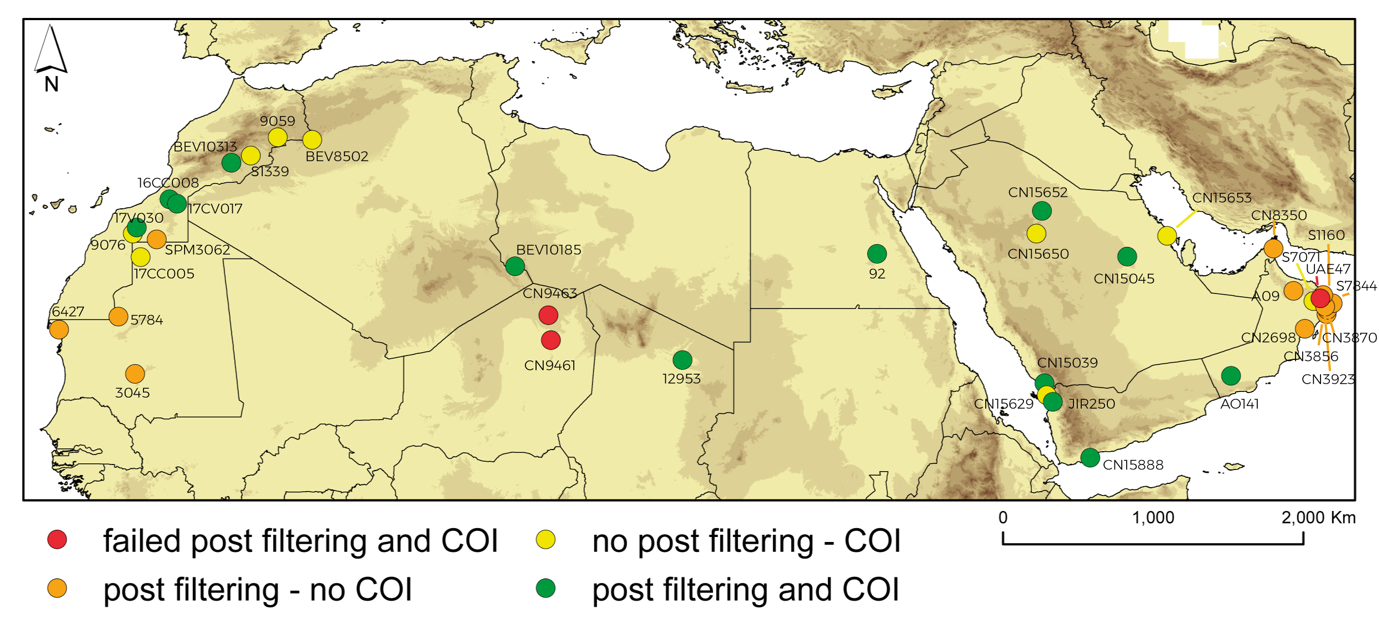

**Fig. S1:** Distribution for all the samples included in the study with colors representing its success or not in both COI and ddRAD sequencing. In red, samples that failed both after the post filtering and COI sequencing, in orange samples that only worked for the ddRADseq post filtering, in yellow samples that only worked for the COI sequencing and in green samples that worked both for ddRADseq and COI sequencing.

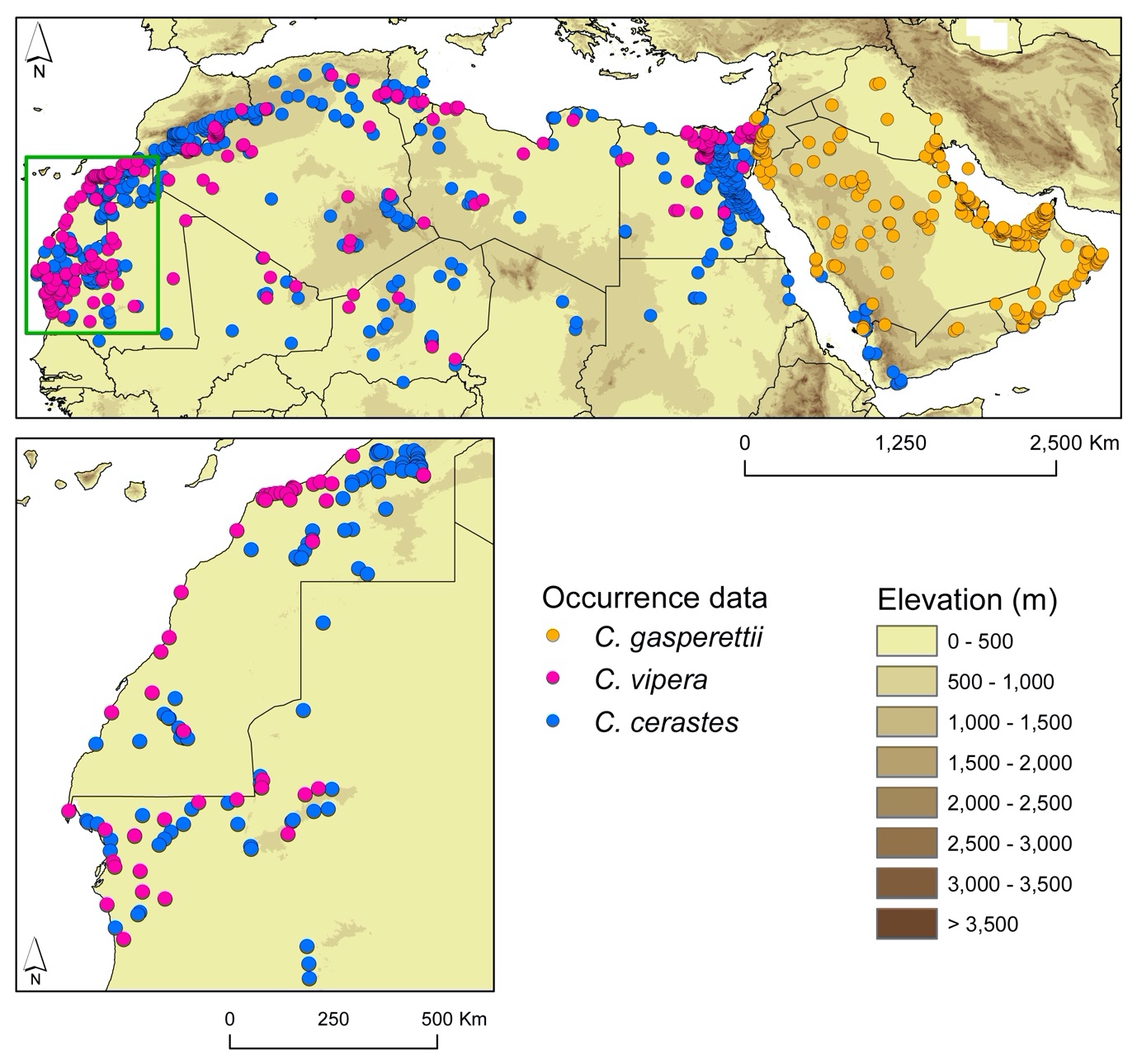

**Fig. S2:** Distribution of occurrence data gathered for the three *Cerastes* species in accordance to the global (top) and regional (bottom) approaches.

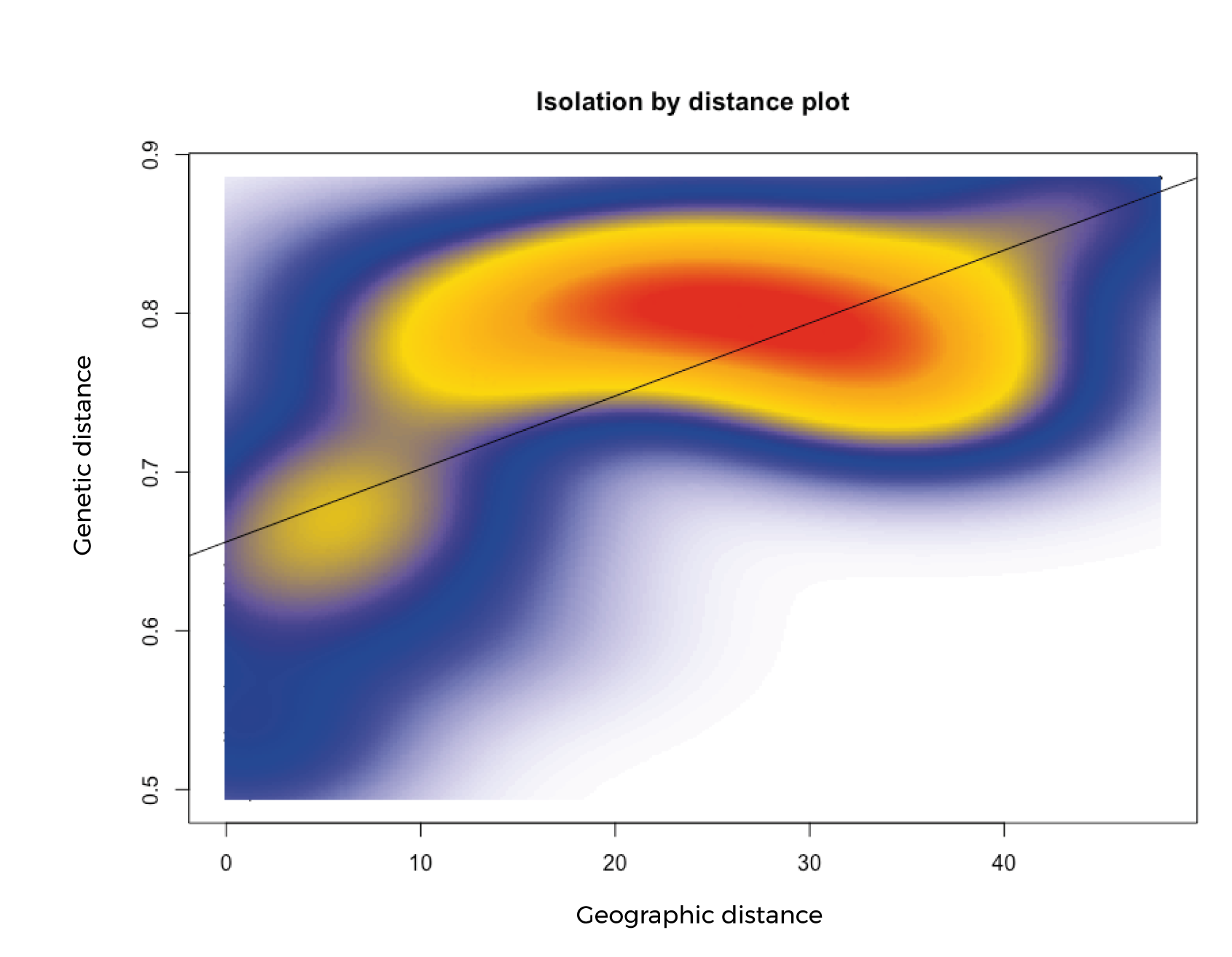

**Fig. S3:** Plot between genetic and geographic distances among all the genomic samples of *Cerastes cerastes*. (r = 0.69, *p*-value < 0.001).

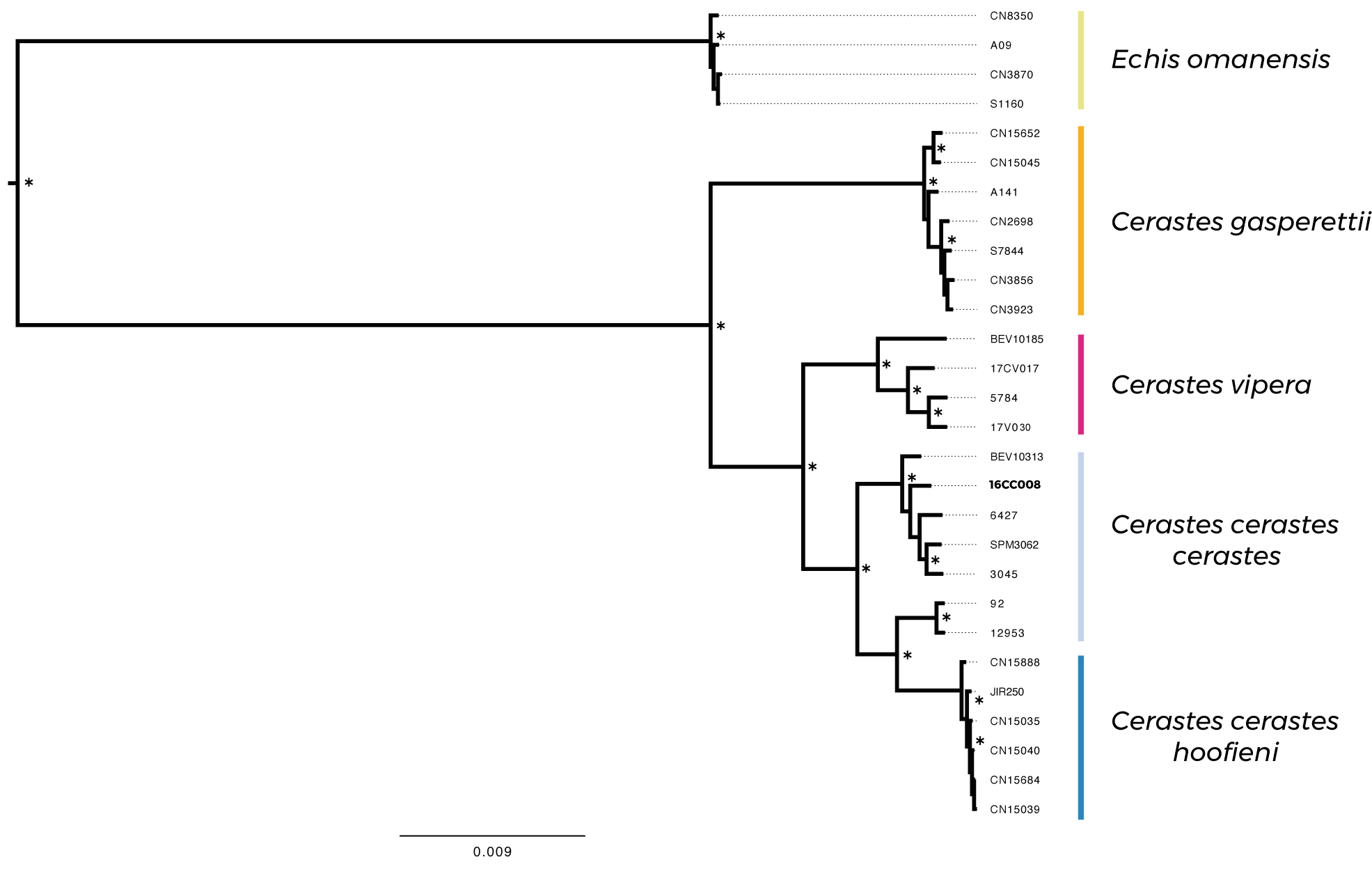

**Fig. S4:** Maximum Likelihood phylogeny reconstructed with 531,260 bp. Nodes with bootstrap higher than 95 are shown with an asterisk. In bold is the putative *C. c.* *mutila* sample.

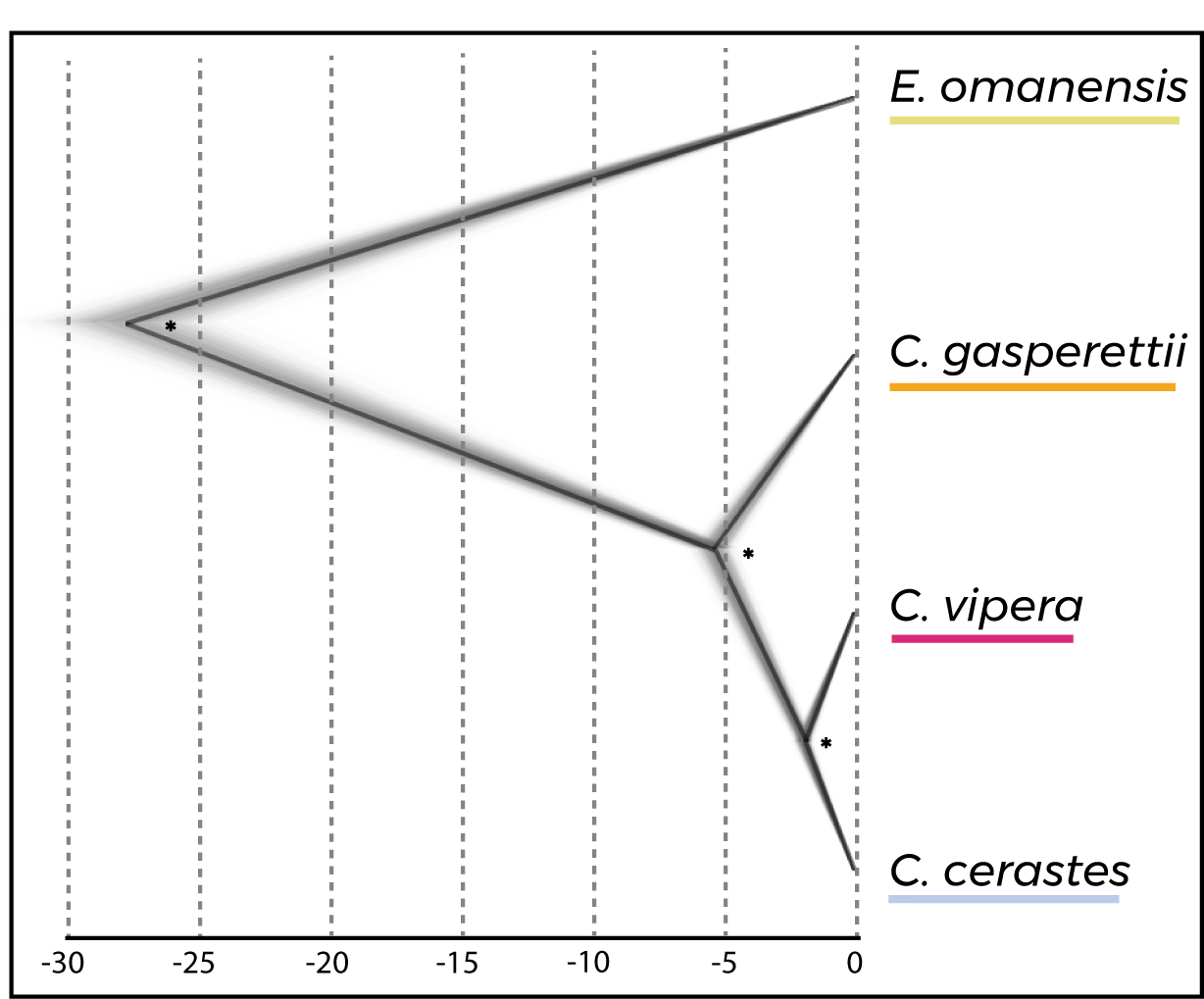

**Fig. S5:** Species tree implemented with SNAPP, including two samples per species over a total of 6,668 *SNPs*. Consensus tree is shown in black while posterior trees are in grey. Posterior probabilities above 0.95 are indicated with an asterisk.

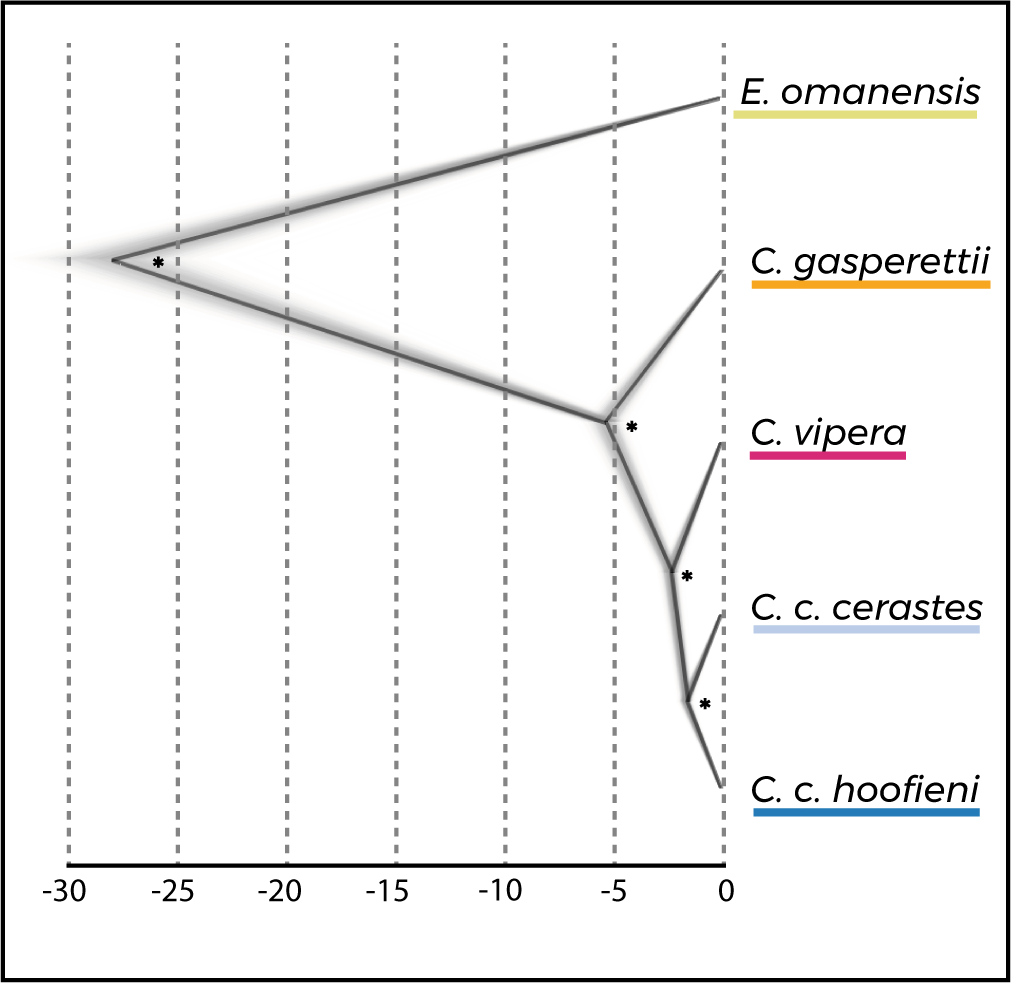

**Fig. S6:** Species tree implemented with SNAPP, including two samples per species, ncluding two individuals of *C. c. hooefini* over a total of 6,378 *SNPs*. Consensus tree is shown in black while posterior trees are in grey. Posterior probabilities above 0.95 are indicated with an asterisk.

**Table S1:** Table with all the samples included in the genomic analyses with information regarding country, coordinates in decimal degrees, ddRADseq raw and filtered reads, sample inclusion after the filtering steps and if COI fragment was amplified. If COI fragment was already available, accession number is shown. Samples used for the species tree analysis (SNAPP) are shown in bold. Accession numbers with asterisk were downloaded from NCBI.

| **Species** | **Code** | **Country** | **Latitude** | **Longitude** | **Raw reads** | **Filtered reads** | **Post filtering** | **COI** |
| --- | --- | --- | --- | --- | --- | --- | --- | --- |
| *C. c. cerastes* | 16CC008 | Morocco | 28.56 | -9.79 | 3,559,557 | 3,552,021 | YES | OR327691 |
| *C. c. cerastes* | 9059 | Morocco | 32.28 | -3.32 | 3,270 | 3,262 | NO | OR327692 |
| *C. c. cerastes* | BEV8502 | Morocco | 32.14 | -1.26 | 423,585 | 422,663 | NO | OR327693 |
| *C. c. cerastes* | S1339 | Morocco | 31.19 | -4.92 | 424,601 | 423,914 | NO | OR327694 |
| *C. c. cerastes* | 13693 | Mauritania | 25.09 | -11.52 | 57,647 | 57,485 | NO | ON943580* |
| *C. c. cerastes* | **3045** | Mauritania | 18.08 | -11.86 | 2,081,481 | 2,077,163 | YES | NO |
| *C. c. cerastes* | 6427 | Mauritania | 20.74 | -16.41 | 899,235 | 897,657 | YES | NO |
| *C. c. cerastes* | CN9461 | Mauritania | 20.11 | 13.03 | 87,802 | 87,682 | NO | NO |
| *C. c. cerastes* | CN9463 | Mauritania | 21.61 | 12.88 | 175,213 | 174,866 | NO | NO |
| *C. c. cerastes* | **BEV10313** | Morocco | 30.75 | -6.09 | 2,203,278 | 2,198,466 | YES | OR327695 |
| *C. c. cerastes* | SPM3062 | Western Sahara | 26.16 | -10.56 | 358,156 | 357,408 | YES | NO |
| *C. c. cerastes* | 9076 | Western Sahara | 26.51 | -11.99 | 217,971 | 217,618 | NO | OR327696 |
| *C. c. cerastes* | 92 | Egypt | 25.29 | 32.55 | 910,543 | 908,663 | YES | OR327697 |
| *C. c. cerastes* | 12953 | Chad | 18.91 | 20.91 | 547,385 | 546,438 | YES | ON943571* |
| *C. c. hoofieni* | CN15035 | Saudi Arabia | 17.52 | 42.58 | 3,954,015 | 3,946,963 | YES | OR327698 |
| *C. c. hoofieni* | CN15039 | Saudi Arabia | 17.52 | 42.58 | 6,144,685 | 6,133,281 | YES | OR327699 |
| *C. c. hoofieni* | CN15040 | Saudi Arabia | 17.52 | 42.58 | 3,264,257 | 3,259,035 | YES | OR327700 |
| *C. c. hoofieni* | CN15684 | Saudi Arabia | 17.52 | 42.58 | 1,865,649 | 1,862,059 | YES | OR327701 |
| *C. c. hoofieni* | CN15629 | Saudi Arabia | 16.80 | 42.70 | 519,464 | 518,641 | NO | OR327702 |
| *C. c. hoofieni* | CN15888 | Yemen | 13.05 | 45.32 | 1,547,767 | 1,545,270 | YES | OR327703 |
| *C. c. hoofieni* | JIR250 | Yemen | 16.40 | 43.07 | 7,897,975 | 7,888,149 | YES | OR327704 |
| *C. vipera* | **17CV017** | Morocco | 28.29 | -9.34 | 1,724,912 | 1,722,092 | YES | OR327705 |
| *C. vipera* | 17V030 | Western Sahara | 26.87 | -11.75 | 1,946,622 | 1,941,611 | YES | OR327706 |
| *C. vipera* | **5784** | Mauritania | 21.52 | -12.85 | 836,709 | 835,078 | YES | NO |
| *C. vipera* | BEV10185 | Algeria | 24.55 | 10.89 | 2,960,609 | 2,954,302 | YES | OR327707 |
| *C. g. gasperettii* | AO141 | Oman | 17.96 | 53.74 | 1,089,989 | 1,085,610 | YES | OR327708 |
| *C. g. gasperettii* | CN15045 | Saudi Arabia | 25.13 | 47.52 | 8,923,691 | 8,908,855 | YES | OR327709 |
| *C. g. gasperettii* | CN15650 | Saudi Arabia | 26.50 | 42.09 | 307 | 300 | NO | OR327710 |
| *C. g. gasperettii* | CN15651 | Saudi Arabia | 27.87 | 42.42 | 104,242 | 104,005 | NO | OR327711 |
| *C. g. gasperettii* | **CN15652** | Saudi Arabia | 27.87 | 42.42 | 436,347 | 434,676 | YES | OR327712 |
| *C. g. gasperettii* | CN15653 | Saudi Arabia | 26.37 | 49.91 | 18,155 | 18,012 | NO | OR327713 |
| *C. g. gasperettii* | CN2698 | Oman | 20.79 | 58.15 | 1,571,647 | 1,566,777 | YES | NO |
| *C. g. gasperettii* | CN3856 | Oman | 21.67 | 59.45 | 5,358,375 | 5,341,541 | YES | NO |
| *C. g. gasperettii* | CN3923 | Oman | 21.94 | 59.50 | 1,083,460 | 1,079,698 | YES | NO |
| *C. g. gasperettii* | S7071 | Oman | 22.45 | 58.67 | 492 | 491 | NO | OR327714 |
| *C. g. gasperettii* | **S7844** | Oman | 22.31 | 59.81 | 3,984,961 | 3,972,856 | YES | NO |
| *E. omanensis* | **A09** | Oman | 23.06 | 57.47 | 1,954,758 | 1,949,128 | YES | NO |
| *E. omanensis* | CN3870 | Oman | 22.11 | 59.36 | 3,778,828 | 3,772,621 | YES | NO |
| *E. omanensis* | **CN8350** | Oman | 25.61 | 56.27 | 2,404,763 | 2,400,950 | YES | NO |
| *E. omanensis* | S1160 | Oman | 22.81 | 59.25 | 4,537,060 | 4,528,532 | YES | NO |
| *E. omanensis* | UAE47 | Oman | 22.62 | 59.09 | 1,107 | 1,103 | NO | NO |
| *E. carinatus* | MG699966 | Afghanistan | - | - | - | - | - | MG699966* |
| *E. carinatus* | MG699965 | Afghanistan | - | - | - | - | - | MG699965* |

**Table S2:** Variables considered in niche overlap analyses

**Global-climate variables** (Worldclim version 2.1 - www.worldclim.org)

BIO3 = Isothermality

BIO6 = Min Temperature of Coldest Month

BIO7 = Temperature Annual Range

BIO10 = Mean Temperature of Warmest Quarter

BIO15 = Precipitation Seasonality (Coefficient of Variation)

BIO16 = Precipitation of Wettest Quarter

BIO17 = Precipitation of Driest Quarter

**Global-landcover variables** (ESA GlobCover ver 2.2 - Bicheron et al., 2008)

rocks

sand

sparse vegetated areas

**Regional-landcover variables** (Campos et al., 2018)

white dunes

vegetated dunes

compact sand

compact and rocky soil

gravel floodplains

gravel floodplains with sand

rocky plateaus + bare rock

**Table S3:** F_st_ (above) and *p-values* (below) at the subspecies level. Significant *p-values* are shown in bold.

|  | *C. c. cerastes* | *C. c. hoofieni* | *C. vipera* | *C. gasperettii* | *E. omanensis* |
| --- | --- | --- | --- | --- | --- |
| *C. c. cerastes* |  | 0.607 | 0.597 | 0.796 | 0.981 |
| *C. c. hoofieni* | **0.001** |  | 0.807 | 0.906 | 0.994 |
| *C. vipera* | **0.004** | **0.003** |  | 0.819 | 0.982 |
| *C. gasperettii* | **0.001** | **0.001** | **0.002** |  | 0.988 |
| *E. omanensis* | **0.003** | **0.004** | **0.031** | **0.003** |  |

**Table S4:** Genetic diversity for all the species. We calculated the parameters for the species *Cerastes cerastes* as well as for the two different subspecies separately.

| **Population** | **Ho** | **Hs** | **Gis** |
| --- | --- | --- | --- |
| *C. cerastes* | 0.015 | 0.045 | 0.675 |
| *C. c. cerastes* | 0.024 | 0.043 | 0.448 |
| *C. c. hoofieni* | 0.005 | 0.006 | 0.221 |
| *C. vipera* | 0.032 | 0.053 | 0.4 |
| *C. gasperettii* | 0.016 | 0.023 | 0.295 |
| *E. omanensis* | 0.013 | 0.018 | 0.256 |

**Table S5**: Quantification of ecological niches and their overlap for the main clades found in *Cerastes* in accordance to three main approaches (global-climate, global-landcover and regional-landcover). It is depicted the variance explained by the three first components of each PCA (Var. PCA), the volumes of the ecological niches for the first (vol x) and second (vol y) clade in the comparison, the volume intersection and union, the unique volumes for each clade and the values of Sørenson (K) and Overlap (OI) indexes. In the latter metrics, bold values mean that statistical test were significant (p < 0.05).

| **dataset** | **comparison (x vs y)** | **Var. three PCs, PCA** | **vol x** | **vol y** | **intersection** | | **union** | **unique x** | **unique y** | **K** | **OI** |
| --- | --- | --- | --- | --- | --- | --- | --- | --- | --- | --- | --- |
| global-climate | *C. gasperettii* vs *C. cerastes*+*C. vipera* | 0.41, 0.224, 0.176 | 67.543 | 152.404 | 53.390 | 166.557 | | 14.153 | 99.014 | **0.494** | **0.800** |
|  | *C. cerastes* vs *C. vipera* |  | 152.354 | 165.558 | 129.013 | 188.899 | | 23.342 | 36.545 | **0.808** | **0.836** |
| global-landcover | *C. gasperettii* vs *C. cerastes*+*C. vipera* | 0.465, 0.341, 0.194 | 42.372 | 58.403 | 32.519 | 68.256 | | 9.853 | 25.884 | **0.649** | **0.765** |
|  | *C. cerastes* vs *C. vipera* |  | 43.869 | 45.516 | 36.342 | 53.043 | | 7.527 | 9.174 | 0.814 | **0.835** |
| regional-landcover | *C. cerastes* vs *C. vipera* | 0.220, 0.204, 0.176 | 91.370 | 78.934 | 59.262 | 111.041 | | 32.108 | 19.672 | **0.662** | **0.709** |
